# Structural insights into type I and type II Lamassu anti-phage systems

**DOI:** 10.1101/2025.06.23.660979

**Authors:** Ming Li, Xiaolong Zhao, Xingyu Zhao, Dong Li, Weijia Xiong, Zirui Gao, Ling Huang, Linfeng An, Yongxiang Gao, Shanshan Li, Yue Feng, Kaiming Zhang, Yi Zhang

## Abstract

Bacteria have developed a variety of immune systems to combat phage infections. The Lamassu system is a prokaryotic immune system with a core conserved Structural Maintenance of Chromosomes (SMC) superfamily protein LmuB and diverse effectors named LmuA, whose mechanism remains unclear. Here, we present a series of cryo-electron microscopy structures of type I Lamassu complex from *Bacillus cellulasensis* and type II Lamassu complex from *Vibrio cholerae,* both in apo- and dsDNA-bound states, which reveal an unexpected stoichiometry and topological architecture, distinct from canonical SMC complexes. Combined structural and biochemical analyses show how the nuclease effector LmuA is sequestered in an inactive monomeric form within the Lamassu complex, and upon sensing foreign DNA ends, dissociates and assembles into an active tetramer capable of DNA cleavage. Our findings elucidate the mechanism by which Lamassu systems detect viral replication and implement anti-phage defense, highlighting the unprecedented roles of SMC proteins in prokaryotic immunity.

---

During the arms race between bacteria and their viruses, bacteria have developed various defense systems to protect against phage infection ^1^. These defense systems function through many different mechanisms, such as degradation of phage nucleic acids, which is utilized by CRISPR-Cas systems, restriction-modification (RM) systems, Argonaute systems, etc ^2-5^. Moreover, recently more and more bacterial defense systems have been found to defend against invading phages via abortive infection (Abi) ^6,7^, a process which involves premature death or growth arrest of the infected cell, thereby preventing phage replication and spread to nearby cells^7^. It has been shown that over 70% of sequenced prokaryotes employ Abi as part of their defensive strategies^6^. Defense systems using Abi include cyclic oligonucleotide-based antiphage signaling systems (CBASS), Thoeris, type III CRISPR-Cas system, toxin-anti-toxin and many others^8-14^. Among the systems via Abi, detection of phage infection will ultimately activate cell-killing protein domains (or effector proteins) through many different mechanisms. Once activated, these domains will result in DNA or RNA degradation, translational arrest, NAD^+^ depletion, ATP degradation, membrane disruption, etc^6^.

Lamassu anti-phage system was first reported in 2018, with two identified components, an endonuclease effector LmuA and a putative DNA sensor, LmuB ^15^. Afterwards, a short gene of unknown function, *lmuC*, was discovered in a subset of Lamassu systems ^16^, which are named as type II Lamassu, therefore the *lmuA-lmuB* two-gene systems as type I Lamassu. The type II Lamassu was also denoted as ATP-binding cassette-three component systems ‘ABC-3C’ ^17^. Later, Lamassu system has been found to be highly diverse, with the endonuclease domain of LmuA replaced by many other effector domains, including SIR2, hydrolase, protease, phosphoesterase, monooxygenase domains, et al. ^18^. Some of these domains have been repeatedly identified in effector proteins of Abi systems ^19^. Notably, type II Lamassu from the El Tor *Vibrio cholerae* strain, was also denoted as DNA defense module DdmABC, with DdmA as LmuA protein, DdmC as LmuB protein and DdmB as LmuC protein ^20^. As the most studied Lamassu system, it has been found to be triggered by palindromic sequences from phages and plasmids ^21^.

Among the components of the system, LmuB belongs to the structural maintenance of chromosomes (SMC) family, a conserved family of ATPases critical for high-order chromosome organization in both prokaryotes and eukaryotes, essential for chromosome maintenance and DNA repair across domains of life ^22,23^. It has been predicted to be structurally similar to the well-studied eukaryotic Rad50 and bacterial SbcC-like proteins ^21^, also known as 3C-ABC ^17^. Both BcLmuA of type I Lamassu from *Bacillus cellulasensis* and VcLmuA comprises an N-terminal Cap4 nuclease domain ^24^ (previously known as DUF4297), their C-terminal domain (CTD) belongs to CTD12 and CTD7 motif, respectively, named according to the ABC-3C systems ^17^. VcLmuC has been predicted to fold as a small globular protein with no predicted homologs, comprising the middle component motif (MC3) in the ABC-3C systems ^17^.

Despite predictions about individual components, the overall architecture, subunit stoichiometry, and activation mechanism of Lamassu systems remain unknown. In particular, how the system senses phage DNA and triggers abortive infection, and how type I and type II variants have evolved and differ in structure and function. Here, we present structural and biochemical insights into both type I and type II Lamassu systems, revealing a conserved activation mechanism in which the effector LmuA is sequestered in an inactive state and released to form an active tetramer upon DNA end sensing. Our findings provide structural and mechanistic insights into how SMC-based Lamassu complexes mediate anti-phage defense.

## Type I and type II Lamassu systems both form heterooligomers

We select type I Lamassu (LmuAB) from *Bacillus cellulasensis* NIO-1130 (short ‘BcLamassu’ hereafter) and type II Lamassu (LmuACB) from *Vibrio cholerae* O1 El (short ‘VcLamassu’ hereafter) for further characterization (Fig.1a). For BcLamassu, initial attempts to purify the components individually revealed that BcLmuB was well-behaved in size-exclusion chromatography (SEC), whereas BcLmuA exhibited severe aggregation (Extended Data Fig.1a,b). Next, co-expression of BcLmuA with BcLmuB enabled successful purification of a stable heterooligomeric complex (Extended Data Fig.1c). SEC coupled with multi-angle light scattering (SEC-MALS) analysis indicated molecular masses of 131.8 ±1.02 kDa for BcLmuB and 165.6 ±1.60 kDa for the BcLmuAB complex, consistent with a LmuB homodimer bound to a single LmuA monomer (Extended Data Fig.1d and Extended Data Fig.1e).

We next turned to the VcLamassu system. Similar to BcLmuA, VcLmuA also showed a tendency to aggregate when expressed alone (Extended Data Fig.1f). VcLmuC was stable and soluble by itself, but VcLmuB was poorly expressed and could not be purified individually (Extended Data Fig.1g,h). After testing various co-expression strategies with different vector combinations and tag configurations, we successfully reconstituted a stable VcLmuACB complex by co-expressing His₆-tagged VcLmuA and untagged VcLmuB from one vector, along with untagged VcLmuC from a second vector (Extended Data Fig.1i). SEC-MALS determined the molecular mass of the purified complex to be 202.7 ±4.79 kDa, consistent with a LmuB homodimer associated with one LmuA and one LmuC monomer (Extended Data Fig.1d and Extended Data Fig.1e). Taken together, these results demonstrate that both type I and type II Lamassu systems assemble into heterooligomeric complexes centered on a LmuB dimer, which serves as a structural core for binding either LmuA alone (type I) or both LmuA and LmuC (type II).

## Lamassu system is activated by DNA ends and regulated by ATP hydrolysis

Interestingly, both BcLmuB and VcLmuB comprise a naturally deviant Walker B motif in their ATPase heads (Fig.1a), raising the question whether they could hydrolyze ATP and how their ATPase activity regulates the anti-phage activity. ATPase assay showed that both BcLmuAB and VcLmuACB do hydrolyze ATP at a moderate rate, similar as Smc-ScpAB from *Bacillus subtilis* reported in a recent study^25^, and this activity was almost abolished by their mutation of Walker A motif residue BcLmuAB^K37A^ or VcLmuACB^K40A^ (Fig.1b). Interestingly, incubation with a 59-bp dsDNA only weakly stimulated the ATPase activity of the WT but not mutated BcLmuAB/VcLmuACB complex, consistent with canonical SMC complexes to some extent, whose ATPase activity can be activated by DNA binding ^26^. In turn, we tested whether the dsDNA binding activity can be regulated by ATP. To eliminate the effects of nuclease activity, we performed DNA-binding analyses using nuclease-dead mutants of two complexes (BcLmuA^K59A^B and VcLmuA^K57G^CB). Electrophoretic mobility shift assay (EMSA) demonstrated that both BcLmuA^K59A^B and VcLmuA^K57G^CB bind to dsDNA (Extended Data Fig.2a), whose binding is enhanced by low concentration of ATP but inhibited by high concentration of ATP (Fig.1c), suggesting that moderate ATP hydrolysis promotes DNA binding but too much ATP saturation at ATPase heads of LmuB may hinder DNA association.

Since the Lamassu complex has been proved to be an active ATPase and it binds to dsDNA, we moved on to test how ATP hydrolysis and dsDNA binding regulates its nuclease activity. First, we tested the *in vitro* nuclease activity of BcLmuAB/VcLmuACB in the presence or absence of ATP. Interestingly, the results showed that, analogous to dsDNA binding, the activity of VcLmuACB is also stimulated by low concentration of ATP (≤1 mM), but inhibited by higher concentration of ATP (≥2 mM), even abolished by 3 mM ATP (Fig.1d and Extended Data Fig.2b). Moreover, the same regulation of VcLmuACB was not observed when ATP was replaced by its slowly hydrolysable form ATPγS (Figure 1d), suggesting that ATP hydrolysis is essential for this regulation. This led us to test whether the nuclease activity of VcLmuACB^K40A^ Walker A motif mutant would also be regulated by ATP, as it loses ATP hydrolysis activity (Fig.1b). Interestingly, this mutant exhibits a rather low nuclease activity under the same conditions as WT protein, and almost barely responds to ATP concentration (Fig.1e), suggesting that ATP binding ability is also essential for activation of the Lamassu system. Results of the BcLmuAB system are similar except that its nuclease activity is not evidently enhanced by low concentration of ATP (Fig.1d,e). Furthermore, adding a 59-bp dsDNA enhanced the nuclease activity mediated pUC19 degradation by both BcLmuAB and VcLmuACB (Fig.1f). Collectively, these findings demonstrate that Lamassu complexes act as ATP-hydrolysis-regulated nucleases, which are activated by the presence of dsDNA ends and tightly controlled by nucleotide concentration.

**Fig. 1.**
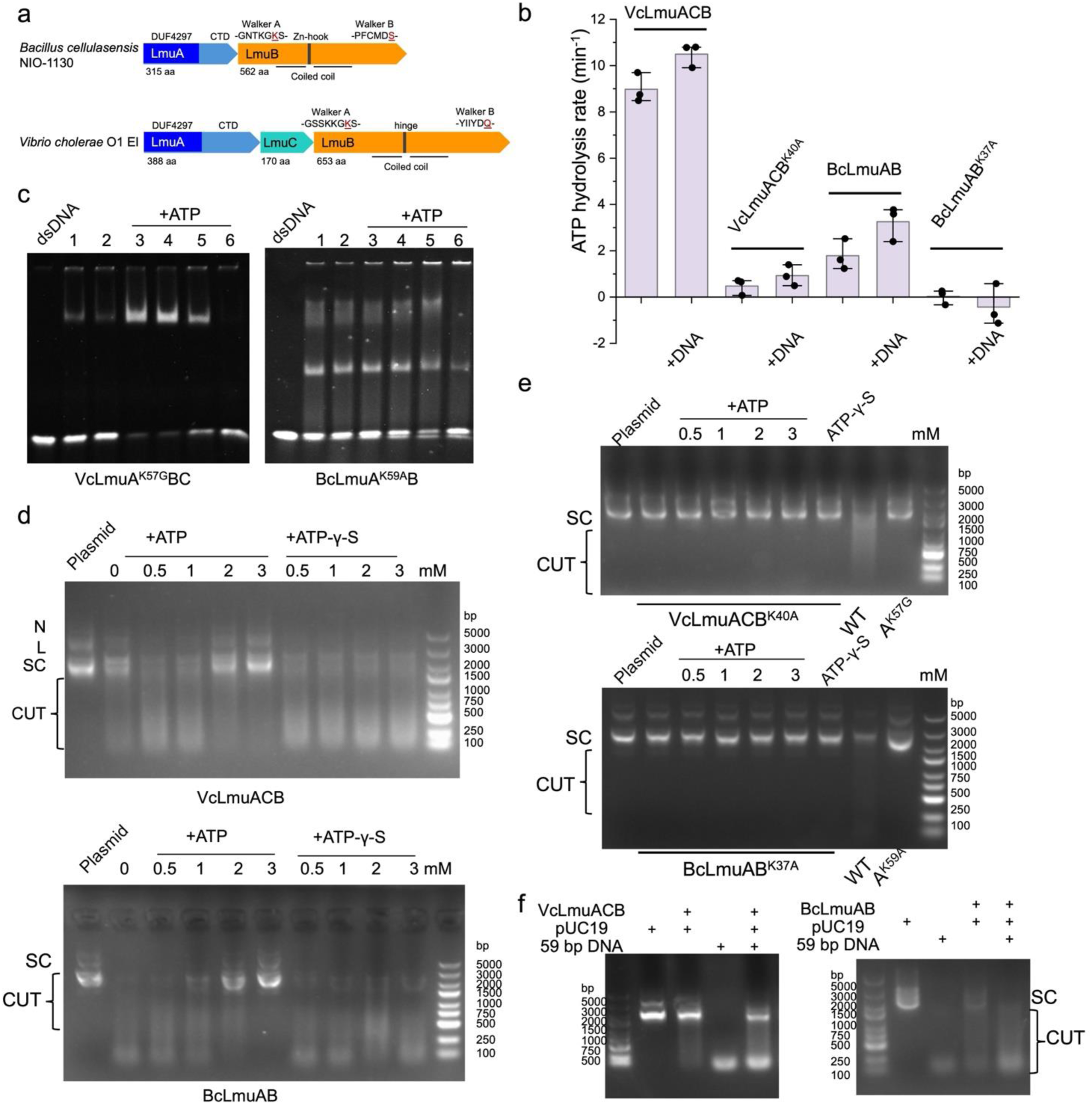
Type I and type II Lamassu complex is activated by DNA ends and regulated by ATP. **a**, Domain architecture of proteins of type I and type II Lamassu systems in this study. Conserved domains and features are shown in the schematics. **b**, Measurement of ATP hydrolysis activity for VcLmuACB, BcLmuAB and variants at 2 μM final with and without 2 μM 59 bp DNA duplex, 1 mM ATP. Each group with three replicate measurements. Means and standard deviations are shown. **c**, EMSA showing that DNA binding by BcLmuAB and VcLmuACB is regulated by ATP. ATP is added at 0.5, 1, 2, 3 mM, respectively, following the order of the triangle. In lanes marked 2-6, 5 mM Mg^2+^ is included. **d**, Agarose gel analysis showed that the nuclease activity of VcLmuACB is regulated by ATP concentration, whereas varying concentrations of the slowly-hydrolyzable analog ATPγS had no significant effect. According to the order indicated in the figure, each lane was supplemented with 0.5, 1,2 or 3 mM ATP or ATPγS, respectively. N: nicked. L: linear. SC: supercoiled. **e**, DNA degradation assays using VcLmuACB mutants demonstrated that the nuclease activity of the ATP-binding site mutant (K40A) was no longer responsive to either ATP or ATPγS concentration. According to the order indicated in the figure, each lane was supplemented with 0.5,1,2, or 3 mM ATP, respectively. ATPγS was added at 3 mM. **f**, DNA degradation assays in the presence of 59-bp dsDNA. BcLmuAB/VcLmuACB complex, pUC19 and 59-bp dsDNA are added with a concentration of 20, 11.4 and 200 nM, respectively.

## Overall structures of the apo type I and type II Lamassu complexes

To gain structural insights into the Lamassu systems, we performed cryo-electron microscopy (cryo-EM) on both the type I (BcLmuAB) and type II (VcLmuACB) complexes in their apo states. The structure of apo BcLmuAB was solved at a resolution of 3.71 Å with the same stoichiometry as determined by SEC-MALS, with BcLmuB: BcLmuA at 2:1 (Fig.2a, Extended Data Fig.3 and Extended Data Table1). Both BcLmuA and the two BcLumB protomers were resolved as full-length proteins. In the structure, two protomers of BcLmuB (BcLmuBα and BcLmuBβ) adopt similar compact conformations, and associate together via tight coiled coil and Zn-hook domain interactions (Fig. 2a). Notably, their globular fold ATPase head domains are separated apart from each other. The coiled coli domain of both BcLmuB protomers forms a kink in the middle position (residues 210-225 and 351-357) (Fig.2a) and two protomers also interact with each other at the kink region. A canonical Zn-hook motif (²⁸⁵CQYC²⁸⁸) of the two LmuB protomers was closely juxtaposed at the most distal end from the ATPase head region (Fig. 2a). However, the local density map lacked sufficient resolution to confirm the presence of Zn atoms. Interestingly, the kink in LmuB creates spatial proximity between the distal region of BcLmuBβ coiled coli domain and BcLmuBα ATPase head (Fig.2a), potentially generating steric hindrance that prevents the assembly of the two ATPase head domain of the two protomers. Contrary to typical SMC systems where accessory proteins bind to the ATPase head domain, BcLmuAB structure reveals a single full-length BcLmuA anchored via its CTD binding to the distal end of the BcLmuB coiled coil domain, forming extensive interfaces with both protomers (Fig.2a). The N-terminal nuclease effector domain of BcLmuA extends out from the complex core. As this domain has been known to require dimerization for DNA cleavage activity ^24^, the monomeric BcLmuA in BcLmuAB complex suggests that the apo-state might represent a resting conformation waiting for activation. This spatial arrangement of LmuA also suggests a potential mechanism for different effector recognition through conformational flexibility of the LmuB arms.

**Fig. 2.**
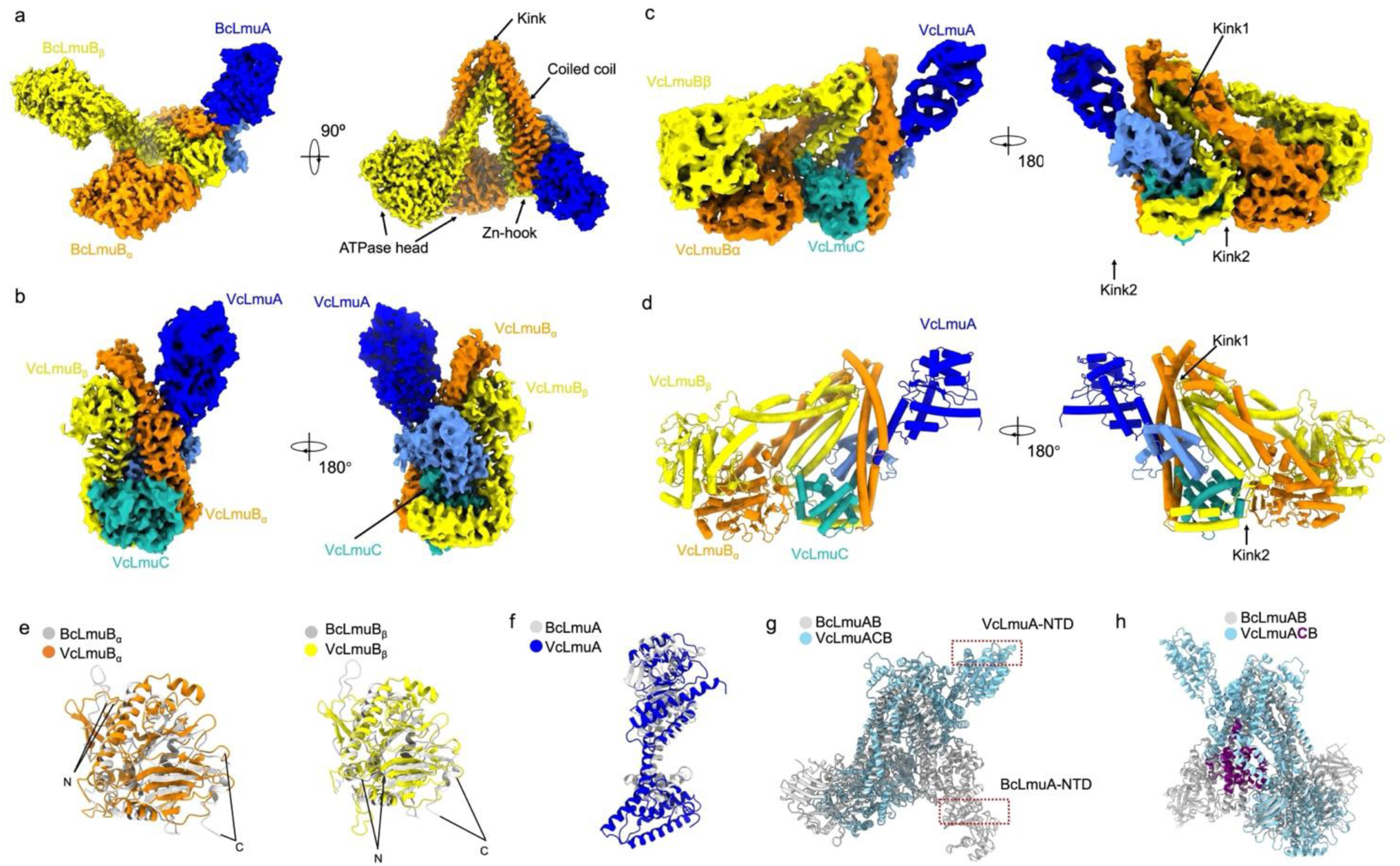
Apo-state structure of type I and type II Lamassu complex. **a-c**, Cryo-EM maps of apo BcLmuAB complex (a), apo VcLmuACB with partial structure (b) and apo VcLmuACB with full-length structure (c). **d**, Atomic model of apo VcLmuACB with full-length structure in c. **e-f**, Structural superimposition of LmuB head domain between BcLmuBα and VcLmuBα (e, left), between BcLmuBβ and VcLmuBβ (e, right) and of LmuA between BcLmuAB and VcLmuACB (f). **g-h**,Structural superimposition between apo BcLmuAB and VcLmuACB, highlighting the position of LmuA-NTD (g), and VcLmuC (h), respectively.

For apo VcLmuACB, we have solved two structures. One was solved using the apo VcLmuACB sample at a resolution of 3.54 Å, which contains full-length VcLmuA/VcLmuC subunits but partial structures for the two VcLmuB protomers (Fig.2b, Extended Data Fig.4 and Extended Data Table 2). The second one (4.07 Å resolution) was obtained from the sample of VcLmuACB mixed with dsDNA, since both the apo and dsDNA-bound states (to be described later) are present in this sample (Fig.2c, Extended Data Fig.5 and Extended Data Table2). Despite being at a lower resolution, this structure contains all components as full-length proteins. As the two structures are basically the same in the resolved regions of the first structure (Extended Data Fig.6a), the second one is taken here to introduce the overall structure of VcLmuACB due to its completeness (Fig.2d). Consistent with the SEC-MALS result, VcLmuACB displays a component ratio of VcLmuA: VcLmuB: VcLmuC as 1: 2: 1. While the head domains of VcLmuB protomers are similar to those of BcLmuB protomers (Fig.2e), distinct from the type I Lamassu complex, the two VcLmuB protomers display markedly different conformations in the coiled coil region (Fig.2d). Specifically, while VcLmuBα displays a single kink at the middle position (residues 284-289 and 413-417) similar as BcLmuB, VcLmuBβ shows another kink (kink2, residues 319-323 and 376-386) apart from kink1 in the middle position (residues 271-278 and 416-424) of the coiled coil domain (Fig.2c,d). The two VcLmuB protomers also converge at both kink1 regions and both hinge regions. While VcLmuA adopts a domain organization analogous to BcLmuA (Fig.2f), anchoring near the head-distal hinge of the VcLmuB dimer via its CTD, with its NTD extending away from the core complex (Fig,2d, 2e), VcLmuA and BcLmuA differ greatly at both NTD and CTD (Fig.2f). Moreover, in the type II Lamassu complex, VcLmuA-CTD displays a distinct binding mode and its NTD extends to a different orientation relative to the core complex, as compared with BcLmuA in the BcLmuAB complex (Fig.2g). As the component only present in the type II Lamassu complex, the globular-folded VcLmuC is positioned in the center of the VcLmuACB complex, interacting with the VcLmuA CTD, as well as the middle regions of coiled coils of both VcLmuB protomers (Fig.2d,2h). A notable feature of the VcLmuACB structure distinct from BcLmuAB is that the two LmuB head domains are juxtaposed but not separated away from each other (Fig.2a, 2c). While ATP is included during grid preparation, there is no density for it. Considering the central position of VcLmuC in the complex and lack of LmuC in BcLmuAB, we propose that the second kink of VcLmuBβ and the contact between head domains of two LmuB protomers might result from the presence of LmuC subunit. Together, these structures reveal that both type I and type II Lamassu complexes adopt asymmetric heterooligomeric assemblies in their apo states, with LmuA held in a monomeric, potentially inactive conformation.

## Interactions among the subunits are essential for anti-phage activity

To elucidate how subunit interactions contribute to Lamassu function, we examined the structural interfaces in both type I and type II complexes. In the type I BcLmuAB complex (Fig.3a-g), E315 and K314 of BcLmuBβ interact with R87 and D561 of BcLmuBα through electrostatic interactions (Fig. 3b). Hydrogen bond mediated interactions are also formed between polar sidechains of the two LmuB protomers, as shown in Figures 3c and 3d. BcLmuA-CTD interacts with the distal region of coiled coil of both LmuB protomers (Fig.3a). Specifically, Y257, Q258 and N259 of BcLmuA interact with Q254, H289 and Q262 of BcLmuBβ, respectively, through polar interactions (Fig.3e). Moreover, K248, K250 and R252 of BcLmuA also interact with acidic side chain residues of BcLmuBα (Fig.3f). R208 of BcLmuA-NTD interacts with the carbonyl oxygen of K291 of BcLmuBα (Fig.3g). Mutations of the BcLmuA-LmuB interface residues decreased binding of BcLmuA to BcLmuB (Fig.3h). Then, we moved on to test the impact of mutating these residues on the anti-phage activity of BcLamassu. First, we confirmed that heterologous expression of BcLmuAB in *Escherichia coli* BL21 demonstrated obvious defense against phage T1 among the *E. coli*-specific phages preserved in our laboratory, which was decreased by catalytic site mutation (K59A) on BcLmuA (Fig.3i). Mutations of the residues in the interface between LmuB protomers and those between LmuA and LmuB also markedly decreased the defense (Fig.3i), suggesting that these interactions are essential for the anti-phage activity of the BcLamassu system.

**Fig. 3.**
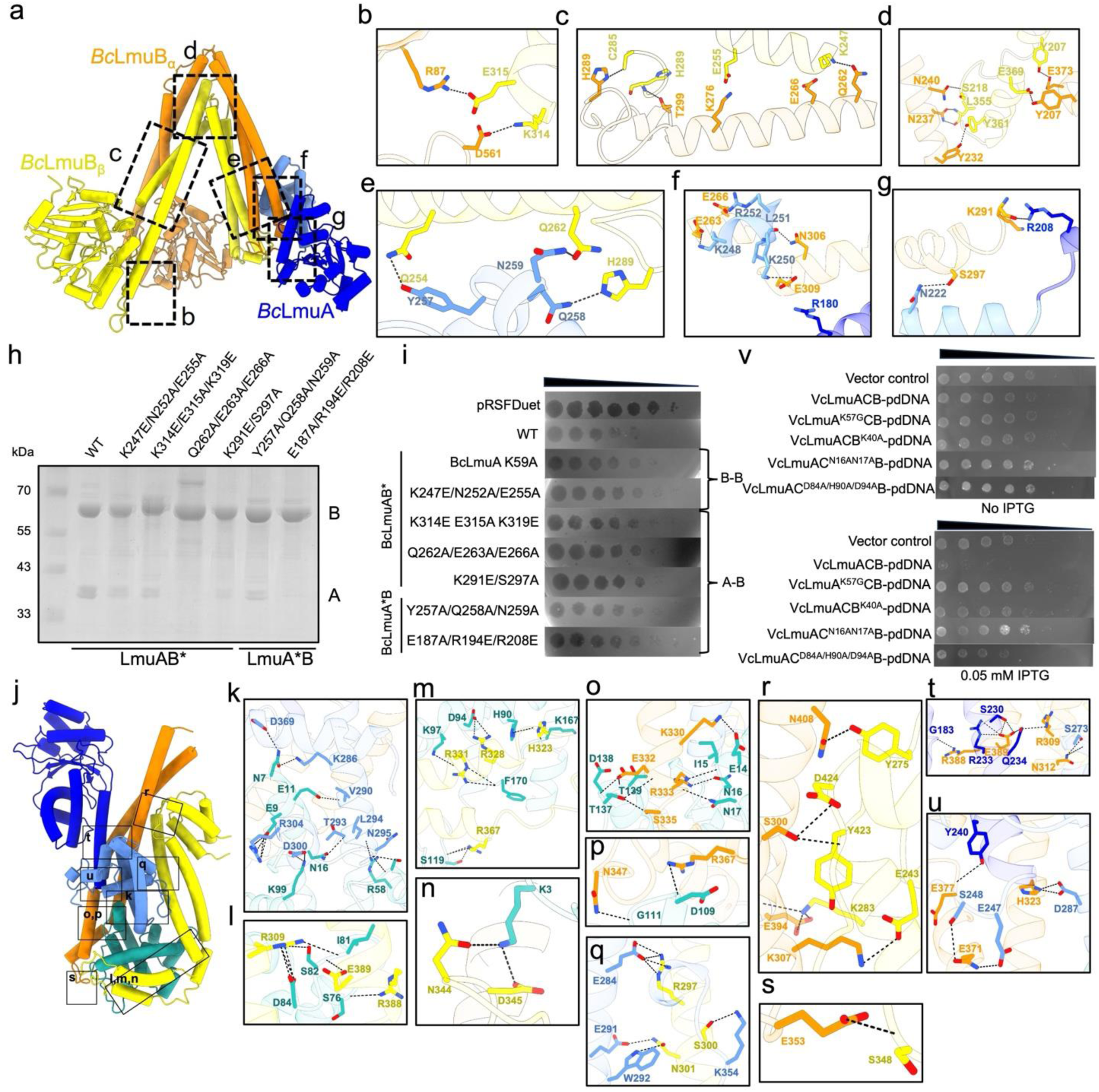
Interface interactions are essential for activity of type I and type II Lamassu system. **a-g**, Atomic model of the BcLmuAB complex is shown in a, in which detailed interactions are shown in b-g. **h**, Purified BcLmuAB WT and mutant complexes were subjected to SDS-PAGE gel, showing the binding ratio between BcLmuB and BcLmuA, in which BcLmuB was N-terminally His-tagged. **l**, Validation of the interface residues of BcLmuAB by plaque assay. **i-u**, Atomic model of the VcLmuACB complex is shown in a, in which detailed interactions are shown in k-s. **v**, Cells encoding the indicated plasmids were plated in 10-fold serial dilution on lysogeny broth (LB)- agar in conditions without induction or induce expression.

Regarding the interface of the type II VcLmuACB complex, we will mainly refer to the one with partial LmuB structures due to its relatively higher resolution. Detailed interactions among LmuA, LmuC and the visible parts of LmuB in the structure are shown in Figures 3j-u. Additional interactions between the two VcLmuB protomers based on the full-length structure are shown in Extended Data Figure 7. Here, we will focus on the interactions between the type II Lamassu characteristic subunit VcLmuC and the other subunits. In the complex, VcLmuC forms extensive interactions with LmuA-CTD and the middle regions of both LmuB protomers. In the VcLmuC-LmuA interface, N7, E11, N16, T55 and R58 of VcLmuC mainly interact with the α helix of VcLmuA containing residues 286-295 (Fig.3k). In the interface between VcLmuC and VcLmuBβ, R309, R388 and E389 of VcLmuBβ interact with the mainchain atoms of I81, S76 and S82 of VcLumC, as well as VcLmuC-D84 sidechain (Fig.3l). VcLmuC-K3 and D94 also interacts with VcLmuBβ-D345 and R328 through electrostatic interaction, respectively (Fig.3m,n). In the VcLmuC-LmuBα interface, E14, N16, N17 of VcLmuC interact with K330 and R333 of VcLmuBα, and N347 and R367 of VcLmuBα also form hydrogen bonds with mainchain atoms of D109 and G111 of VcLmuC (Fig.3o). Next, we moved on to test the effects of mutations of these interface residues on the activity of VcLmuACB. No obvious anti-phage activity was observed through plasmid-based protein expression under our experimental conditions, which might be why *E. coli* strains carrying the LmuACB transposons integrated into the chromosome were used to show anti-phage activity in the previous study ^20^. Then we co-expressed VcLmuACB with palindromic DNA (short for “pdDNA” hereafter) fragment, after induction, *E. coli* cultures exhibited decreased bacterial growth (Fig.3v) as reported in a recent study ^21^. The pdDNAs are predicted to form stem-loop hairpins from single-stranded DNA, which may trigger cell density-dependent death (CDD) dependent on VcLmuACB. Consistently, both Walker A motif disruption (K40A) on VcLmuB and nuclease catalytic site mutation (K57A) on VcLmuA substantially attenuated VcLmuACB-mediated bacterial toxicity (Fig.3v), suggesting that this effect stems from VcLamassu itself rather than protein overexpression burden or plasmid maintenance stress. Similarly, mutations of the VcLmuC interface residue N16/N17 or D84A/H90A/D94 markedly alleviated the toxicity by co-expression of VcLmuACB and pdDNA (Fig.3v). Taken together, these results illustrate the structural and functional importance of the binding interface residues of both type I and type II Lamassu systems in antiphage defense and CDD of cells.

## Mechanism of DNA-end sensing by Lamassu complex

To understand the mechanism of activation of Lamassu by phage DNA, we moved on to solve its structures complexed by dsDNA. To this end, both type I and type II Lamassu complex in this study was mixed with a 59-bp dsDNA sample, respectively, supplemented by ATP/Mg^2+^. Similarly, we will first introduce the type I BcLmuAB-DNA complex (3.73 Å resolution), which revealed a complex of BcLmuAB_2_ bound to one end of a resolved 41-bp dsDNA molecule (Fig.4a, Extended Data Fig.8 and Extended Data Table 1). Interestingly, structural alignment of the apo- and dsDNA-bound BcLamassu showed that BcLamassu does not exhibit marked conformational changes upon DNA binding, with an RMSD of 1.165 Å for 1281 Cα atoms (Extended Data Fig.6b). The head domain of the two BcLmuB protomers are still away from each other (Fig.4a). Slight shifts are observed for both BcLmuB, especially in the head and coiled coil region near the head domain. Surprisingly, BcLamassu specifically binds the dsDNA end, instead of binding DNA using the ATP-engaged SMC heads as in canonical SMC-DNA complexes. The dsDNA mainly interacts with the head domain of BcLmuBα using one of its ends, as well as the LmuA-NTD using its middle region (Fig. 4b-d). Mutations of the DNA interacting residues of the BcLamassu complex markedly decreased the defense (Fig.4e), suggesting that these interactions are essential for the anti-phage activity of the BcLamassu system. This suggests that the Lamassu complex possibly detects DNA ends by direct binding, instead of sliding onto the DNA as proposed for other SMC complexes. While the middle region of dsDNA is stabilized by the LmuA-NTD, we propose that it represents a sensing-competent but catalytically inactive state. This is supported by two observations: (1) BcLmuA remains monomeric, and (2) its catalytic residue (K59) does not contact the DNA substrate (Fig.4d). Thus, DNA-end binding may serve as a trigger for activation, preceding effector domain dimerization and nuclease engagement.

**Fig. 4.**
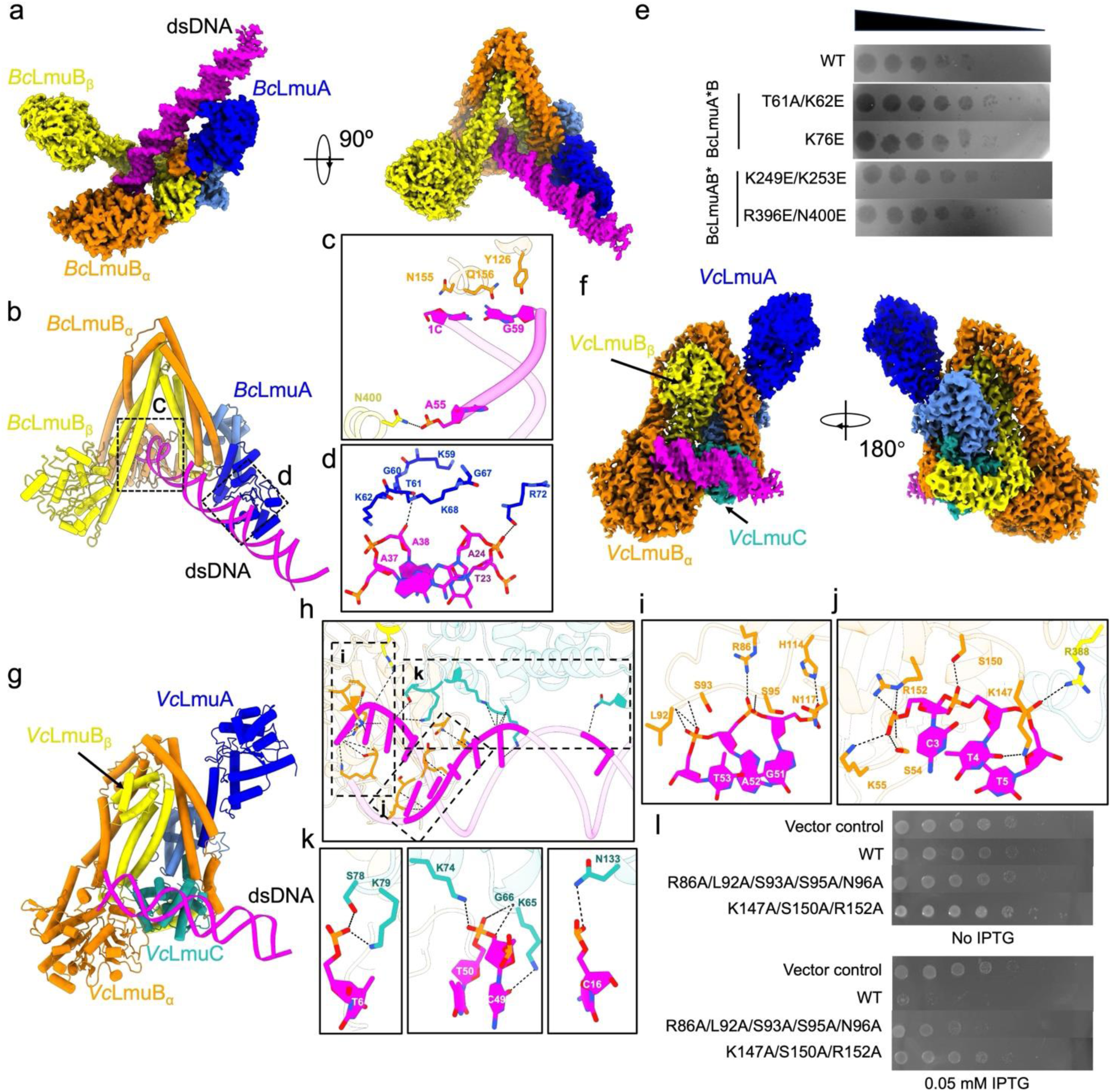
DNA-bound state structure of type I and type II Lamassu complex. **a**, Cryo-EM map of BcLmuAB complexed with dsDNA. **b-d**, Model of the BcLmuAB-dsDNA complex is shown in b, in which detailed interactions between dsDNA and BcLmuAB are shown in c-d. **e**, Validation of the DNA binding residues of BcLmuAB by plaque assay. **f**, Cryo-EM map of VcLmuACB complexed with dsDNA. **g-k**, Atomic model of the VcLmuACB-dsDNA complex is shown in g, in which detailed interactions between dsDNA and BcLmuAB are shown in h-k. **l**, Cells encoding the indicated plasmids were plated in 10-fold serial dilution on lysogeny broth (LB)- agar in conditions without induction or induce expression. WT or LmuB mutated VcLmuACB is tested.

Structure of the type II VcLamassu-DNA complex was solved at 2.95 Å resolution, which revealed a complex of VcLmuACB_2_ bound to a 27-bp dsDNA fragment (Fig.4f,Extended Data Fig.9 and Extended Data Table 2). In the structure, density for the head and part of coiled coil region that links to the head domain of VcLmuBβ are missing, possibly due to its flexibility, which also suggests that the two head domains of the VcLmuB protomers are not engaged together despite inclusion of ATP and Mg^2+^. Similar as type I Lamassu, structural alignment of the apo- and dsDNA-bound VcLamassu also revealed no apparent conformational changes between the two structures, with an RMSD of 0.754 Å among 1180 Cα atoms (Extended Data Fig 6c,d). Type II VcLamassu also binds the dsDNA ends partially with the VcLmuBα head domain as type I BcLamassu, however, the rest of dsDNA is stabilized by VcLmuC but not VcLmuA-NTD (Figure 4G), consistent with the fact that VcLmuA-NTD extends towards a distinct orientation from BcLmuA-NTD relative to the core complex. Progression of the dsDNA is blocked by the head domain of VcLmuBα, mainly from a loop spanning residues 129-152 (Fig.4h). Together, this suggests that VcLamassu also senses dsDNA ends as BcLamassu complex. Analysis of the binding interface of DNA in the VcLamassu complex shows that R86, H114, N117 of VcLmuBα interact with phosphate groups of one strand of dsDNA through electrostatic interactions (Fig.4i). Moreover, the phosphate groups also polarly interacts with mainchain atoms of L92, S93 and S95 of VcLmuBα. K55, K147 and R152 of VcLmuBα interact with phosphate groups of the other strand of dsDNA through electrostatic interactions (Fig.4j). In addition, R388 of VcLmuBβ also interacts with a phosphate group of the strand, forming the only interaction from VcLmuBβ. In the VcLmuC-DNA interface, S78, K65, K74, K79 and N133 of VcLmuC interact with phosphate groups of the dsDNA (Fig.4k). Consistently, mutation of the DNA interacting residues of the Lamassu complex also alleviated the toxicity by co-expression of VcLmuACB and pdDNA (Fig. 4l). Taken together, both type I and type II Lamassu complexes sense ends of dsDNA to be activated, which is markedly distinct from previously characterized SMC-DNA complexes.

## Structural basis of activated LmuA

Having established the resting and DNA-sensing states of both type I and type II Lamassu systems, we next investigated how the LmuA effector becomes activated. Intriguingly, in the VcLmuACB samples incubated with dsDNA and ATP/Mg²⁺, a distinct particle population emerged, corresponding to a tetrameric assembly of VcLmuA (Fig.5a,b, Extended Data Fig.9 and Extended Data Table 2). In the structure, VcLmuA tetramerizes through its CTD and extends their NTD out (Fig.5b). While it has been known that Cap4 nuclease domain dimerizes to be activated, interestingly, VcLmuA tetramer does not form two but only one active site in the middle. For simplicity, the four protomers in the structure are named VcLmuAα, β, γ and δ, respectively, according to the clockwise order viewed from top of the complex (Fig.5a,b). Among the protomers, the NTD of VcLmuAα and γ extends out nearly parallel to their CTD and interlocks with each other to form a V-shaped dimer, displaying a predicted DNA degradation pocket. However, the NTD of VcLmuAβ and δ extends almost vertically to their CTD. No obvious conformational changes are observed in the CTD among the four LmuA protomers and that in the VcLmuACB complex (Fig.5c). However, a striking structural variation occurs to the region linking CTD to NTD, spanning residues 234-259, which displays three different conformations among the five LmuA molecules. In the LmuA of the VcLmuACB complex, this region forms a loop (residues 246-259) and a helix (residues 234-245) which connects to the helix (residues 219-233) to form a long helix (Fig.5d). However, in the four VcLmuA protomers, this region forms a loop-helix-loop conformation and extends towards a markedly different orientation compared to that in the VcLmuACB complex. Notably, the loop spanning residues 234-238 of the four LmuA protomers also extends in two almost vertical orientations, with LmuA α and γ in the same conformation, and LmuAβ and δ in the same conformation. When the NTD of the five LmuA molecules is aligned together, marked conformational changes are observed, also with LmuA α and γ in the same conformation, and LmuAβ and δ in the same conformation (Fig.5e,f). Importantly, the NTD of LmuAβ/δ is more structurally similar to that in the VcLmuACB complex, again suggesting that VcLmuA α/γ forms an active dimer in the LmuA tetramer (Fig.5f). Notably, severe steric clashes can be observed when VcLmuA tetramer is aligned to VcLmuACB complex on LmuA (Extended Data Fig.10a,b), again suggesting that LmuA is kept in an inactivated conformation in the Lamassu complex. Such a significant conformational switch especially in VcLmuA α/γ might be facilitated by a complete disengagement of LmuA from the Lamassu complex upon activation. Since no DNA density is observed in the LmuA tetramer, we made a structural model of LmuA-NTD-DNA complex with AlphaFold3 ^27^, which overlaps well with the NTD dimer of VcLmuAα and γ (Fig.5g). This again suggests that LmuA tetramer represents an activated form of LmuA. Multiple interactions can be found between VcLmuA-CTDs and between VcLmuA-NTDs. Detailed and representative interactions between NTDs of VcLmuAα and γ, NTDs of VcLmuAα and β, CTDs of VcLmuAα and δ, and CTDs of VcLmuA γ and δ are shown in Figures S10C-S10M. Mutants of the VcLmuA CTD interface residue R312A and R312E both abolished the toxicity by co-expression of VcLmuACB and pdDNA (Fig.5h).

Inspired by this finding of VcLamassu system and to capture DNA in the LmuA tetramer, next we incubated BcLmuAB (BcLmuA K57A mutation to inactive the nuclease domain) with dsDNA plus ATP/Mg^2+^ and subjected it to cryo-EM studies. Again, particle populations corresponding to BcLmuA tetramer as well as BcLmuB-dsDNA are observed (Extended Data Fig.11). However, unfortunately, due to relatively low resolution (5.16 Å for BcLmuA tetramer and 6.53 Å for BcLmuB-dsDNA), we could not build the atomic models. Then we made a tetramer model of BcLmuA with Alphafold3, which also revealed a similar conformation as the VcLmuA tetramer (Extended Data Fig.12a). Notably, mutation of the residues in the modeled tetramer interface of BcLmuA N239A/T240A/I240A also decreased the anti-phage activity of BcLamassu (Fig. 5i and Extended Data Fig.12), suggesting that LmuA adopts the same mechanism to be activated in type I and type II Lamassu systems. Together, in the presence of dsDNA plus ATP/Mg^2+^, LmuA transitions from a monomeric, inactive effector in the resting Lamassu complex to an active tetrameric nuclease to cleave DNA.

**Fig. 5.**
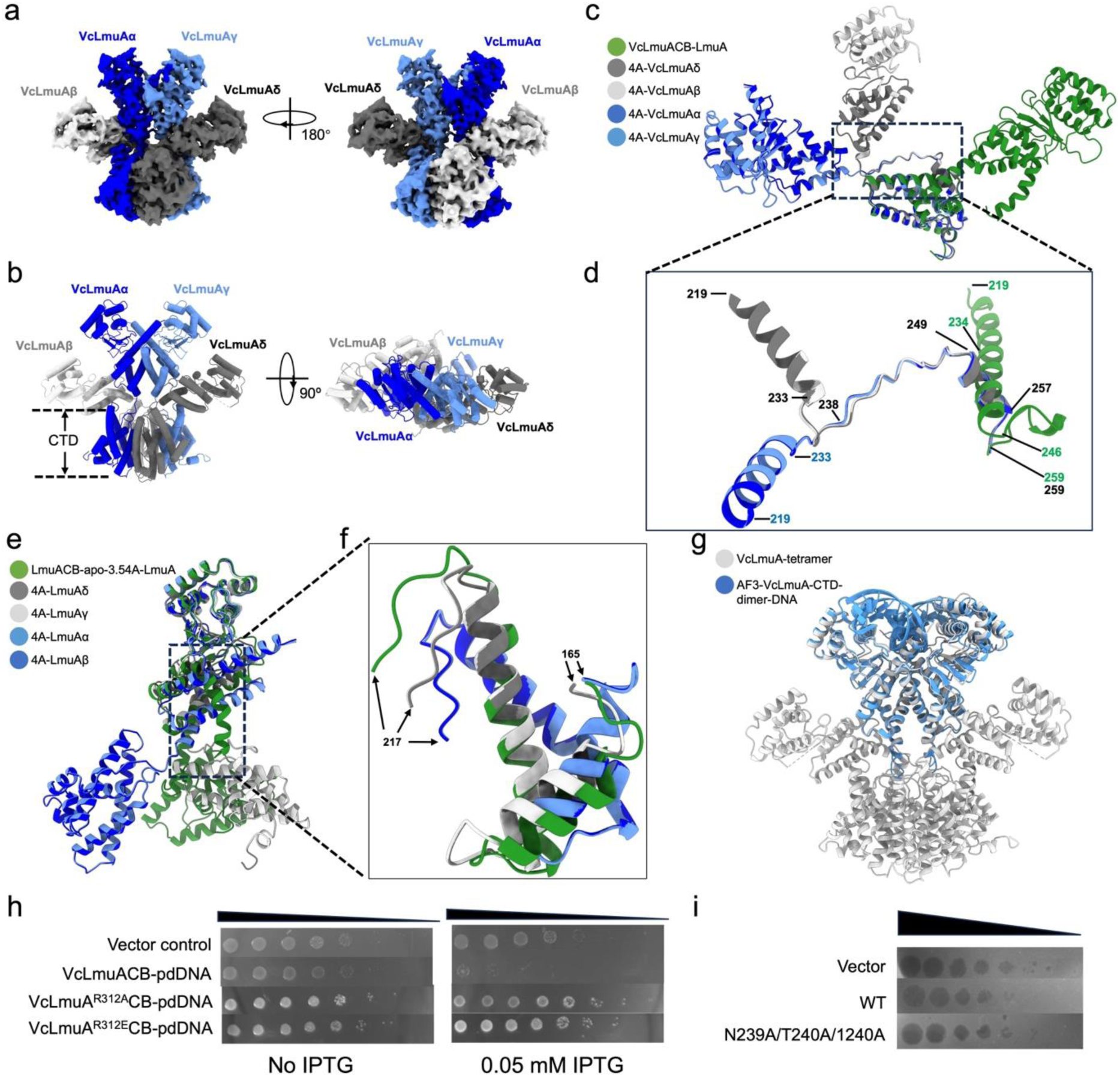
Structural basis of the activated form of LmuA. **a**, Cryo-EM map of VcLmuA tetramer with C2 symmetry. **b,** Atomic model of VcLmuA tetramer. The CTD region is marked. **c-d**,The CTDs of the four VcLmuA protomers in the tetramer are aligned with VcLmuA-CTD of the VcLmuACB apo state (c). The region with the most remarkable structural difference is shown in D. **e-f**, The NTDs of the four VcLmuA protomers in the tetramer are aligned with LmuA-NTD of the VcLmuACB apo state. The region with the most remarkable structural difference is shown in f. **g**, The AlphaFold3 (AF3) predicted complex structure of VcLmuA-CTD dimer with dsDNA is aligned with the VcLmuA tetramer. The dsDNA is aligned in the DNA cleavage site of the CTD dimer in the middle of the VcLmuA tetramer. **h**, Validation of the tetramer interface residues of BcLmuA by pdDNA-induced cell death assay. **i**, Validation of the predicted tetramer interface residues of BcLmuA by plaque assay.

## Discussion

Prokaryotes have evolved multiple anti-phage defense systems to trigger abortive infection, thereby inhibiting the phage propagation. Here, our study reveals how an SMC system could be co-opted for phage defense and abortive infection, in which the SMC protein LmuB exhibits multiple unique mechanisms distinct from other canonical SMC complexes. First, rather than forming a symmetrical dimer by SMC proteins, in both type I and type II Lamassu systems LmuBs fold as asymmetrical dimers with different 3D structures from each other. Second, distinct from canonical SMC systems where accessory proteins associate with ATPase heads to regulate hydrolysis ^28^, both LmuA and LmuC localize to coiled coil regions away from ATPase heads. Third, our study implies that Lamassu systems recognize DNA ends by directly binding them, as opposed to clamping and gating function by the coiled coils proposed for related Rad50/Mre11 complexes ^26^. Last but not least, Lamassu system operates through sequestration of LmuA effector under resting state and disengagement of the effector to oligomerize it upon activation.

During preparation of our manuscript, two separate studies also reported mechanisms of type II Lamassu systems through charactering this system from *Salmonella enterica* ^25^and *Vibrio cholerae* ^29^(the same system as our study), respectively. Consistently, their studies also reveal a sequestration state of LmuA in the LmuACB complex, and an activated tetramer conformation of LmuA upon recognition of dsDNA end. Interestingly, Li et al found that DNA binding by SeLmuACB complex is inhibited by ATP, and DNA degradation by LmuACB does not require ATP. However, the other study by Haudiquet et al found that DNA degradation by VcLmuACB complex is enhanced by incubation with ATP and dsDNA. In our study, we have tried a series of concentrations of ATP and found that the nuclease activity of Lamassu system is enhanced under low ATP concentrations but inhibited by high ATP concentrations (Figure 1D). We think that this might reflect a strategy adopted by bacteria to inhibit the accidental activation of the Lamassu system in physiological conditions, considering that the physiological ATP concentration was found to be over 3 mM in *E. coli* ^30^. Interestingly, it has been reported that upon phage infection several bacterial anti-phage systems decrease cellular ATP concentrations through hydrolyzation of ATP^31-33^, which might facilitate activation of Lamassu system under low ATP concentrations.

Our results support a working model of type II Lamassu (the dominant type) as follows (Fig.6), which is also applicable to type I Lamassu. First, the LmuACB complex scans along dsDNA in bacterial cells. Physiological ATP concentration in bacterial cells may stabilize the LmuACB complex. Upon phage infection, free DNA ends or a specific DNA structure might be produced, which facilitates dissociation of the LmuA subunit, meanwhile decrease of ATP concentration by bacterial anti-phage system or phages would further promote disruption of the LmuACB-dsDNA complex and release of LmuA. Then LmuA tetramerizes into an activated state and nonspecifically cleaves cellular DNA. While LmuA could comprise various effector domains, the ubiquitous CTD indicates that disengagement of LmuA from the LmuACB complex for activation and subsequent oligomerization through its CTD might be a conserved mechanism for the activation of different Lamassu systems.

**Fig. 6.**
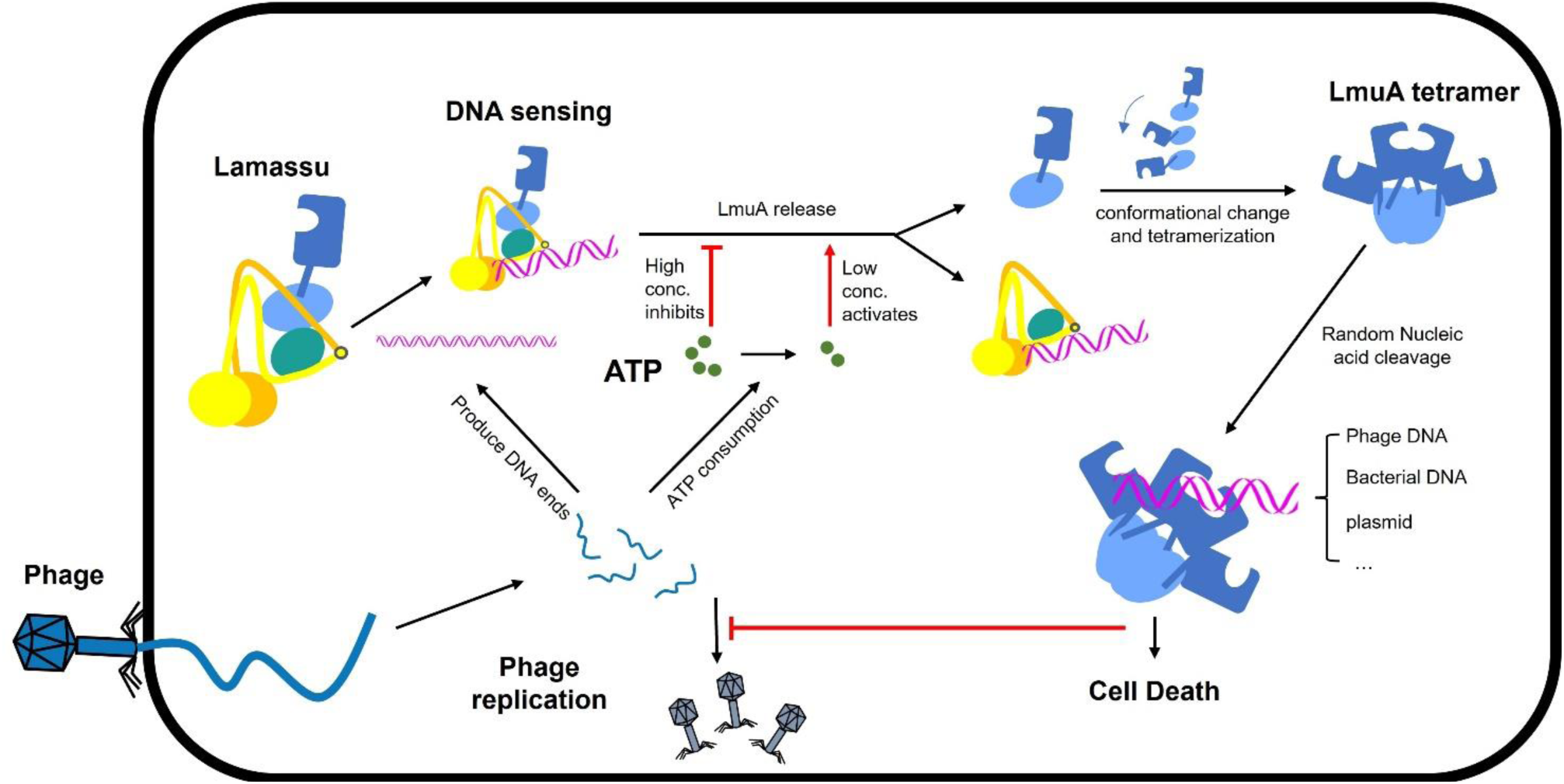
Working model of Lamassu anti-phage system. The native LmuACB complex exists in a LmuA-sequestered state, which enables intracellular DNA scanning and recognition through ATP hydrolysis mediated by LmuB. Following phage invasion, the phage replication may generate abundant free DNA ends or a specific DNA structure that are subsequently sensed by LmuACB complex. Concurrently, phage invasion may directly or indirectly induce ATP consumption via activation of other phage defense systems. Upon stimulation by DNA ends and low ATP concentrations, the LmuACB complex becomes activated. LmuA liberated from the complex, undergoes conformational change, and tetramerizes to form an active non-sequence-specific DNA-cleavage machinery. Consequently, this mechanism induces death of phage-infected cells while blocking phage replication.

Interestingly, the recent preprint by Haudiquet et al also performed bioinformatics studies into Lamassu systems and classified it into short- and long-form Lamassu systems ^29^. Based on their classification, both Lamassu systems in our study belong to the short Lamassu systems, where VcLamassu is in the clade F with Cap4 LmuA, and BcLamassu is in the clade S with loss of LmuC subunit. Notably, our structural and biochemical studies revealed that type I and type II Lamassu systems are different in many aspects apart from composition. In terms of architecture, the ATPase heads of the two LmuB protomers are juxtaposed in VcLmuACB but away from each other in BcLmuAB. They also differ in the binding mode and orientation of LmuA subunit. In terms of function, both nuclease activity and DNA binding of VcLmuACB exhibits obvious enhancement by low concentration of ATP but inhibition by high concentration of ATP. However, BcLmuAB only exhibits inhibition by high concentration of ATP in these two aspects. These differences between type I and type II Lamassu system might reflect their different evolutionary position as investigated by Haudiquet et al ^29^. Overall, our study provides a structural and mechanistic framework for understanding how Lamassu systems utilize an SMC scaffold for immune function. However, several questions remain and need further investigation. First, Lamassu systems feature a variety of effector domains beyond nucleases, raising the question of whether these effectors, such as protease and phosphoesterase domains, share the same oligomerization-based activation mechanism or involve distinct regulatory strategies. Second, it remains unclear how phage infection precisely triggers LmuA release. Do specific DNA-end structures or protein–DNA intermediates act as signals, and is there any crosstalk with other bacterial stress or signaling pathways? Another promising avenue is understanding how phages may counteract Lamassu. Given that Lamassu activation is tightly linked to DNA-end sensing and ATP levels, phage-encoded inhibitors might target LmuB’s ATPase activity, mask DNA ends, or hijack the activation switch of LmuA. The interplay between Lamassu and other cellular systems, such as energy depletion modules or DNA repair pathways, could also be explored to better understand abortive infection as a holistic response.

## Methods

### Cloning and Protein Purification

For type-I Lamassu, the full length LmuA and LmuB gene from *Bacillus cellulasensis* NIO-1130 were synthesized by GenScript and codon-optimized for expression in *E. coli*. They were amplified by PCR and clone into a pRSFDuet vector with LmuB in multiple cloning site 1 and LmuA in multiple cloning site 2. LmuB was fused to a 6×His-tag, while LmuA was expressed without a tag. The mutants were generated by rolling circle amplification and were overexpressed and purified in the same way as for the WT protein. All of the protiens were transformed into *E.coli* strain BL21 (DE3) grown in SB medium at 37℃, until the OD600 up to 1.0, then the temperature was shift to 16℃, and 0.3 mM IPTG was added to induce protein expression for 16-20 h, then the cell pellet was harvested by centrifugation and resuspended in lysis buffer (50 mM Tris-HCl pH 8.0, 500 mM NaCl, 10 mM Imidazole, 5% glycerol,1 mM DTT), and lysed by sonication. The supernatant was harvested by centrifugation for 1h at 10,000 RPM, and load into 3 ml Ni-NTA Excel Sepharose (Cytiva) pre-equilibrated with lysis buffer, then the sepharose was washed by 30 ml lysis buffer50 mM Tris-HCl pH 8.0, 500 mM NaCl, 20 mM Imidazole, 5% glycerol, 1mM DTT), then washed with 30 ml lysis buffer supplemented with 20 mM Imidazole and finally eluted with 30 ml lysis buffer supplemented with 400 mM Imidazole. The elution was concentrated to 0.5 ml by 30 kDa cut-off (Milipore)and loaded into a Superdex-200 increase 10/300 column (Cytiva) pre-equilibrated with gel filtration buffer (20 mM Tris-HCl pH 8.0,100 mM NaCl,1 mM DTT). The peak fraction was check by SDS-PAGE and used for cryo-EM sample preparation or 10% glycerol was added and frozen in liquid nitrogen and stored at -80℃ for biochemical assays.

For type-II Lamassu, the full length LmuA, LmuC and LmuB gene from *Vibrio cholerae* O1 El were synthesized by GenScript and codon-optimized for expression in *E. coli*. LmuA and LmuB were amplified by PCR and cloned into a pRSFDuet vector with LmuA in multiple cloning site 1 and LmuB in multiple cloning site 2. LmuA was fused to a 6×His-tag, while LmuB was expressed without a tag. LmuC was amplified by PCR and clone into a pETDuet vector and expressed without a tag. The mutants were generated by rolling circle amplification and were overexpressed and purified in the same way as for the WT protein. All of the proteins were co-transformed into *E. coli* strain BL21 (DE3) and induced by 0.2 mM IPTG when the cell density reached an OD600 of 0.8. After growth at 18 °C for 12 h, the cells were collected, resuspended in Ni-A buffer (50 mM Tris-HCl pH 8.0, 500 mM NaCl, 30 mM imidazole and 5% glycerol) with 1 mM PMSF and lysed by sonication. The cell lysate was centrifuged at 20,000 g for 50 min at 4 °C to remove cell debris. The supernatant was filtered with a 0.45 µm filter and then applied onto HisTrap HP column (Cytiva) and contaminant proteins were removed with Ni-A buffer. The protein was then eluted with a gradient concentration of 0–100% Ni-B buffer (50 mM Tris pH 8.0, 500 mM NaCl, 500 mM imidazole and 5% glycerol). Next, the eluant was desalted into Q-A buffer containing 25 mM Tris pH 8.0,150 mM NaCl, 2 mM DTT and 5% glycerol by a desalting column (Cytiva), and was further purified by ion-exchange chromatography using the HiTrap Q HP column (Cytiva). The protein bound to the column was eluted with a gradient concentration of 150–1000 mM NaCl. The eluant was concentrated and further purified using the Superdex-200 increase 10/300 column (Cytiva) equilibrated with a buffer containing 10 mM Tris–HCl pH 8.0, 500 mM NaCl, 5 mM DTT and 5% glycerol. The purified proteins were analyzed by SDS–PAGE. The fractions containing the target protein complex were pooled, concentrated and stored at - 80℃ for biochemical assays.

### DNA preparation

The 59 bp DNA fragment was commercially synthesized (Sangon Biotech) with both strands purified through preparative polyacrylamide gel electrophoresis (PAGE). Equimolor amounts of the strands were dissolved in SEC buffer (50 mM Tris-HCl pH 8.0,500 mM NaCl,400mM Imidazole,5% glycerol,1mM DTT), subjected to thermal denaturation at 95℃ for 5 min, followed by gradual annealing at room temperature. For pUC19 plasmid preparation, pUC19-harboring *E.coli* (DH5a) was harvested by centrifugation (6,000 rpm,10 min), with plasmid DNA subsequently extracted using Plasmid Extraction Kit (TIANGEN Biotech) according to the manufacturer’s instruction.

### *In vitro* Nuclease Assays

For the cleavage of pUC19, a total of 20 uL reaction contained 20 nM BcLmuAB or VcLmuACB complex and 11.4 nM pUC19 plasmid were suspended in the reaction buffer (20 mM Tris-HCl pH 7.5,100 mM NaCl, 1 mM DTT, 5% glycerol and 5 mM MgCl_2_).The reaction at 37°C for 2 min and quenched by the 2× quenching buffer (95% formamide,1% SDS, 10 mM EDTA, 0.025% (wt./vol.) xylene cyanol, and 0.025% (wt./vol.) bromphenol blue. separated on 1% agarose gels in 1×TBE buffer and stained with Goldview nucleic acid stain, and then visualized by ultraviolet illumination.

### Cryo-EM sample preparation and data collection

For apo-state BcLmuAB, gel filtration eluate was concentrated to 30 μM, centrifuged at 13,000 rpm for 10 min, and 3 μL of the supernatant (supplemented with 0.03% NP-40; Biosharp) was applied to glow-discharged Quantifoil R2/1 200-mesh copper grids. After 5 s incubation, grids were blotted for 3.5 s (blot force 0) at 4 °C and 100% humidity, then plunge-frozen in liquid ethane using a Vitrobot Mark IV (Thermo Fisher Scientific). For the BcLmuAB–dsDNA complex, 10 μM protein was incubated with 10 μM 59-bp dsDNA and 5 mM MgCl₂ on ice for 30 min prior to vitrification. Cryo-EM data for the both samples were acquired using a Titan Krios G3i (Thermo Fisher Scientific) operated at 300 kV and equipped with a K3 Summit direct electron detector (Gatan). Images were recorded using EPU software at 105,000× magnification with a calibrated pixel size of 0.82 Å (Table S1). Exposure time was 3 s with a dose rate of 17.8 e⁻/s/Å².

For the BcLmuA(K59A)B–dsDNA–ATP complex, 6 μM protein was incubated with 10 μM 59-bp dsDNA, 5 mM MgCl₂, and 2 mM ATP at room temperature for 2 h before vitrification. Data were collected using a 200 kV Glacios microscope (Thermo Fisher Scientific) with a Falcon 3 detector (Thermo Fisher Scientific), at 105,000× magnification (pixel size: 1.20 Å), 2 s exposure, and a dose rate of 22.5 e⁻/s/Å² (Extended Data Table1).

For the apo VcLmuACB complex (state 1), 3 μL of 1.5 mg/mL VcLmuACB was gently mixed with 2.04 μL TBS300 (20 mM Tris pH 7.8, 300 mM NaCl), 0.36 μL 1% OG (n-octyl-β-D-glucopyranoside), and 0.6 μL 10× T4 DNA Ligase Buffer (NEB). For state 2, 4.5 μL of VcLmuACB was mixed with 0.2 μL TBS2000 (20 mM Tris pH 7.8, 2 M NaCl), 3.6 μL pUC19 plasmid (233 ng/μL), and 0.4 μL 10× Ligase Buffer, and incubated on ice for 5 min. For the VcLmuACB–dsDNA complex, samples were prepared by mixing 3 μL of VcLmuACB with either 2.4 μL (dataset 1) or 2.22 μL (datasets 2 and 3) of 40 μM 59-bp dsDNA, along with 0.6 μL 10× Ligase Buffer. Datasets 2 and 3 additionally contained 0.18 μL 1% OG. For all VcLmuACB samples, 3 μL aliquots were applied to glow-discharged Quantifoil R2/1 200-mesh gold grids, blotted for 4.5 s, and plunge-frozen in liquid ethane using a Vitrobot Mark IV at 4 °C and 100% humidity. Grids were initially screened on a Glacios (200 kV) microscope and imaged on a 300 kV Titan Krios G3i microscope (Thermo Fisher Scientific) equipped with a K3 Summit detector. Images were collected at 105,000× magnification, with a pixel size of 0.82 Å using EPU software (Extended Data Table2).

### Cryo-EM data processing

Cryo-EM data were processed using standard pipelines in the software packages cryoSPARC^34^ and EMAN2^35^. All dose-fractioned movies were subjected to patch motion correction, patch CTF estimation, particle picking and extraction, 2D classification, ab initio 3D reconstruction, heterogeneous refinement, and non-uniform refinement.

#### BcLmuAB Datasets

For the apo-state BcLmuAB, 2,630 micrographs were selected for template-based particle picking after motion correction and CTF estimation. A total of 1,312,792 particles were extracted with a box size of 366 pixels (bin 3). Following 2D classification and ab initio 3D reconstruction, several rounds of heterogeneous refinement yielded 481,446 selected particles, which were re-extracted with a box size of 336 pixels (bin 1) and subjected to non-uniform refinement. The final reconstruction reached a resolution of 3.71 Å (Extended Data Fig.3).

For the BcLmuAB–dsDNA complex, 12,235 micrographs were processed using the BcLmuAB apo map as a picking template. From these, 1,144,023 particles were extracted (box size 366, bin 3), and after similar processing steps, 437,756 particles were re-extracted (box size 336, bin 1) and refined, resulting in a final map at 3.73 Å resolution (Extended Data Fig.8).

For the LmuB–DNA and LmuA tetramer, 249 micrographs were selected for both blob and template picking, yielding 59,724 particles (box size 256, bin 1). After 2D classification and ab initio reconstruction, 17,938 particles for LmuB–DNA and 28,799 for LmuA tetramer were subjected to non-uniform refinement, resulting in maps at 6.53 Å and 5.16 Å resolution, respectively (Extended Data Fig.11). The workflow described here is depicted in Extended Data3,8 and 11.

#### VcLmuACB Apo and DNA-Bound Datasets

For the apo-state 1 of VcLmuACB, 2,743 micrographs yielded 1,535,228 particles (box size 384, bin 4). After 2D classification, ab initio modeling, and several rounds of heterogeneous refinement, 481,446 particles were re-extracted (box size 336, bin 1) and further refined, resulting in a final map at 3.54 Å resolution.

For apo-state 2 of VcLmuACB, 1,519 micrographs were processed using blob and template picking, yielding 2,360,724 particles (box size 384, bin 4). Following 2D classification and six-class heterogeneous refinement, 180,439 particles (from three classes) were re-extracted (box size 512, bin 1). Further refinement of the sixth class yielded 51,655 particles; another 30,935 were selected from the remaining classes. After reference motion correction a total of 67,762 particles were obtained. Additionally, 23,072 particles with clear DdmC-dimer features from the VcLmuACB–dsDNA dataset were merged into this workflow. Then non-uniform refinement was performed using the combined dataset to resolve the final map at 4.07 Å resolution (Extended Data Fig.5).

#### VcLmuACB–DNA and VcLmuA Tetramer Complexes

For the VcLmuACB–DNA complex, 10,667 micrographs were processed. After motion correction and CTF estimation, particles were extracted with a box size of 384 pixels. Following 2D classification, ab initio 3D reconstruction, and heterogeneous refinement, 164,644 particles (class 1 of 3) were selected and subjected to non-uniform refinement, DeepEMhancer post-processing, reference motion correction, and a second round of non-uniform refinement. The final map reached 2.95 Å resolution and clearly resolved the bound dsDNA (Extended Data Fig.9).

For the VcLmuA tetramer, particles were extracted with a box size of 336 pixels after blob and template picking. After 2D classification, ab initio 3D reconstruction, and heterogeneous refinement with C2 symmetry, 35,249 particles (class 1 of 4) were selected for non-uniform refinement. Following DeepEMhancer post-processing and reference motion correction, a second round of refinement yielded a map at 3.64 Å resolution (Extened Data Fig.9).

### Model building and refinement

The full-length sequence of BcLmuA, BcLmuB, VcLmuA, VcLmuB, and VcLmuC were imported into the AlphaFold3 server^27^ for the initial model generation of their protomers, respectively. The initial models of protomers were first rigidly fitted into the cryo-EM maps using ChimeraX^36^ applying “fit in map tool”, and molecular dynamics flexible fitting (MDFF)^37^ was applied to flexibly fit the atomic model into the map, followed by optimization with Coot^38^ and phenix.real_space_refine^38^. The bound DNA was manually adjusted by Coot and refined by phenix.real_space_refine. For the apo form, the final model of the DNA-bound form was initally fitted into its cryo-EM map, followed by optimization with Coot and phenix.real_space_refine. The bound DNA was manually adjusted by Coot and refined by phenix.real_space_refine. The final models were evaluated by MolProbity^39^. Statistics of the model building are summarized in Extended Data Tables1 and 2. All figures were prepared using Chimera^40^ and ChimeraX.

### ATPase assay

ATPase activity measurements were done with 2 μM final BcLmuAB and VcLmuACB in 150 μL pyruvate kinase/lactate dehydrogenase coupled reactions at 37℃ in ATPase buffer (25 mM Hepes pH 7.5, 50 mM KCl, 5 mM MgCl_2_, 1 mM MnCl_2_, 0.1 mg/mL BSA and 1 mM DTT). The reactions contained 0.5 mM NADH, 2 mM phosphoenol pyruvic acid, 20 U pyruvate kinase, 30 U lactate dehydrogenase, 1 mM ATP and 2 μM 59 bp dsDNA if needed. ADP accumulation was monitored for 20 min by measuring absorbance changes at 340 nm caused by NADH oxidation in an EnSpire Multimode Plate Reader (PerkinElmer). Results were analysed and plotted using the GraphPad Prism 8 software.

### Phage plaque assay

For the plaque assay, the relevant plasmids or their mutants were transformed into *Escherichia coli* strain BL21(DE3). Bacteria containing the defense system and control bacteria lacking the system were incubated overnight at 37 °C in LB medium. A total of 400 µL of bacterial culture was mixed with 5 mL of 0.8% LB-agar supplemented with 0.09 mM IPTG (pre-heated to 65 °C), poured onto the surface of a 2.5% LB-agar plate, and allowed to dry at room temperature for 15 min. Ten-fold serial dilutions of the phages were prepared, and 1.2 µL of each dilution was spotted onto the bacterial layer. After the spots had dried, the plates were inverted and incubated at 37 °C for 3–6 h before imaging.

### Electrophoretic mobility shift assay

Binding between *Bc*LmuA^K59A^B or *Vc*LmuA^K57G^CB complex and the 59 bp dsDNA. 59 bp dsDNA was prepared by annealing the forward and complementary strands in a 1:1 ratio, with the 5′-FAM label at forward strand. Reactions were performed by incubating *Bc*LmuA^K59A^B (0, 0.25, 0.5, 1, 2, 4 and 8 μM) or *Vc*LmuA^K57G^CB (0, 0.5, 1, 2, 4, 8 and 16 μM) complex with 0.25 μM dsDNA for 20 min at 37°C in a 20-μl buffer system containing 20 mM Tris pH7.5, 100 mM NaCl, 1mM DTT, 2 mM ATP and 5 mM Mg^2+^. In order to explore the effect of ATP concentration, reactions were performed by incubating 3 μM *Bc*LmuA^K59A^B or 4 μM *Vc*LmuA^K57G^CB complex with 0.25 μM dsDNA and 0, 0.5, 1, 2, 3 μM ATP for 20 min at 37°C in a 20-μl buffer system containing 20 mM Tris pH7.5, 100 mM NaCl, 1mM DTT and 5 mM Mg^2+^. The mixtures were separated using discontinuous native polyacrylamide gels and visualized by fluorescence imaging.

### Multi-angle light scattering

Multi-angle light scattering experiments were performed in 20 mM Tris pH7.5, 500 mM NaCl and 2 mM DTT using a Superdex-200 10/300 GL size exclusion column (Cytiva). All protein complex concentrations were diluted to 1.0 mg/ml. The chromatography system was connected to a Wyatt DAWN HELEOS laser photometer and a Wyatt Optilab T-rEX differential refractometer. Wyatt ASTRA 7.3.2 software was used for data analysis.

### Cell toxicity assay

pRSFDuet, pETDuet and pACYCDuet (encoding VcLmuA/B, VcLmuC and palindromic DNA, respectively) were transformed into *E. coli* strain BL21(DE3). The cells were first grown to OD_600 nm_ of 1.0. Then 10-fold serial dilutions in LB were performed for each of the samples and 1.2 μL drops were put on agar plates containing 30 μg/mL chloramphenicol, 80 μg/mL ampicillin, 50 μg/mL kanamycin with or without 0.05 mM IPTG. Plates were incubated overnight at 37°C. The negative control (empty vectors) and all of the mutants were analyzed in the same way as above.

Palindromic DNA sequence: CTCAAGGTCATTCGATAAGAGTGGCCTTTATGAA

## Data availability

Cryo-EM structures and atomic models generated in this study have been deposited in the wwPDB OneDep System under EMD accession codes EMD-64575, EMD-64576, EMD-64585, EMD-64586, EMD-64583, EMD-64584 and PDB ID codes under accession codes 9UX7, 9UX8, 9UXK, 9UXL, 9UXH and 9UXI. This paper does not report original code. Any additional information required to reanalyze the data reported in this paper is available from the corresponding authors upon request.

## Acknowledgements

We thank the Cryo-EM Center at the University of Science and Technology of China (USTC) for their assistance with the cryo-EM experiments. This work is supported by National key research and development program of China (2024YFA0916903, 2022YFC3401500, 2022YFA1302700, 2022YFC2303700 and 2022YFC2104800), the National Natural Science Foundation of China (32371329, 32171274, 32371345, 32301044 and 32471301), Scientific Research Innovation Capability Support Project for Young Faculty (ZYGXQNJSKYCXNLZCXM-B1), the Fundamental Research Funds for the Central Universities (QNTD2023-01), Anhui Provincial Natural Science Foundation (2308085QC80), the Strategic Priority Research Program of the Chinese Academy of Sciences (XDB0490000), the Center for Advanced Interdisciplinary Science and Biomedicine of IHM (QYPY20220019), and the USTC Research Funds of the Double First-Class Initiative (YD9100002048 and YD9100002044).

## Author contributions

Y.Z., K.Z. and Y.F. conceived and supervised the project and designed experiments. M.L., X.Z., D.L., W.X., Z.G., L.H. and L.A. purified the proteins and performed *in vitro* activity analysis and in *vivo* assays. M.L., X.Z., Y.G. and K.Z. collected the cryo-EM data and solved the cryo-EM structures. S.L. built and refined the models. Y.Z. wrote the original manuscript. Y.F., K.Z. and Y.Z. revised the manuscript.

