## Extended Data Figures and Table for "Structural insights into type I and type II Lamassu anti-phage systems"

### Extended Data Table 1|Cryo-EM collection, processing and model validation of BcLmuAB datasets

| | BcLmuAB-apo | BcLmuAB-DNA | BcLmuB-DNA | BcLmuA-tetramer |
|---|---|---|---|---|
| **Data collection and processing** | | | | |
| Microscope | Titan Krios G3i | Titan Krios G3i | FEI Glacios | FEI Glacios |
| Voltage (kV) | 300 | 300 | 200 | 200 |
| Camera | Gatan K3 | Gatan K3 | Thermo Falcon3 | Thermo Falcon3 |
| Magnification | 105,000× | 105,000× | 120,000× | 120,000× |
| Pixel size (Å) | 0.82 | 0.82 | 1.2 | 1.2 |
| Total exposure (e-/Å2) | 50.4 | 52.5 | 45 | 45 |
| Exposure time (s) | 3 | 3 | 1.99 | 1.99 |
| Number of frames per exposure | 30 | 30 | 32 | 32 |
| Energy filter slit width (eV) | 20 | 20 | 20 | 20 |
| Data collection software | EPU 2.7 | EPU 2.7 | EPU 2.7 | EPU 2.7 |
| Number of exposures per hole | 4 | 4 | 1 | 1 |
| Defocus range (μm) | -1.6 to -3.1 | -1.6 to -3.1 | -1.8 to -2.8 | -1.8 to -2.8 |
| Number of micrographs collected | 3,194 | 13,330 | 254 | 254 |
| Number of micrographs used | 2,630 | 12,235 | 249 | 249 |
| Number of initial particles | 1,312,792 | 1,144,023 | 59,724 | 59,624 |
| Symmetry | C1 | C1 | C1 | C2 |
| Number of final particles | 447.569 | 437,756 | 17,938 | 28,799 |
| Resolution (0.143 gold standard FSC, Å) | 3.71 | 3.73 | 6.53 | 5.16 |
| **Atomic model refinement** | | | | |
| Software | phenix | phenix | N/A | N/A |
| Clashscores, all atoms | 11.15 | 22.87 | N/A | N/A |
| Poor rotamers (%) | 0 | 0.36 | N/A | N/A |
| Favored rotamers (%) | 99.36 | 99.41 | N/A | N/A |
| Ramachandran outliers (%) | 0.21 | 0.05 | N/A | N/A |
| Ramachandran favored (%) | 95.36 | 95.95 | N/A | N/A |
| MolProbity score | 1.90 | 2.13 | N/A | N/A |
| Bad bonds (%) | 0.00 | 0 | N/A | N/A |
| Bad angeles (%) | 0.03 | 0.03 | N/A | N/A |

**Extended Data Table 2|Cryo-EM collection, processing and model validation of VcLmuACB datasets**

| | VcLmuACB-apo-1 | VcLmuACB-apo-2 | VcLmuACB-DNA complex | VcLmuA tetramer |
|---|---|---|---|---|
| **Data collection and processing** | | | | |
| Microscope | Titan Krios G3i | Titan Krios G3i | Titan Krios G3i | Titan Krios G3i |
| Voltage (kV) | 300 | 300 | 300 | 300 |
| Camera | Gatan K3 | Gatan K3 | Gatan K3 | Gatan K3 |
| Magnification | 105,000× | 105,000× | 105,000× | 105,000× |
| Pixel size (Å) | 0.82 | 0.82 | 0.82 | 0.82 |
| Total exposure (e-/Å2) | 54.4 | 52.8 | 53\52.9\48.3 | 53\52.9\48.3 |
| Exposure time (s) | 3.2 | 2\1.7 | 3.2\3.2\1.7 | 3.2\3.2\1.7 |
| Number of frames per exposure | 32 | 30 | 32\32\30 | 32\32\30 |
| Energy filter slit width (eV) | 20 | 20 | 20 | 20 |
| Data collection software | EPU 2.7 | EPU 2.7 | EPU 2.7 | EPU 2.7 |
| Number of exposures per hole | 4 | 4 | 4 | 4 |
| Defocus range (μm) | -1.5 to -2.9 | -0.9 to -2.9 | -0.9 to -3.1 | -0.9 to -3.1 |
| Number of micrographs collected | 2,788 | 7,758 | 10,939 | 10,939 |
| Number of micrographs used | 2,743 | 7,239 | 10,667 | 10,667 |
| Number of initial particles | 1,535,228 | 10,027,902 | 8,112,757 | 21,632,622 |
| Symmetry | C1 | C1 | C1 | C2 |
| Number of final particles | 122,532 | 78,853 | 164,496 | 30,148 |
| Resolution (0.143 gold standard FSC, Å) | 3.54 | 4.07 | 2.95 | 3.64 |
| **Atomic model refinement** | | | | |
| Software | phenix | phenix | phenix | phenix |
| Clashscores, all atoms | 18.14 | 22.87 | 13.25 | 7.75 |
| Poor rotamers (%) | 0 | 0.36 | 0.39 | 0.23 |
| Favored rotamers (%) | 99.03 | 99.41 | 98.84 | 98.49 |
| Ramachandran outliers (%) | 0.66 | 0.05 | 0.07 | 0 |
| Ramachandran favored (%) | 93.3 | 95.95 | 95.36 | 98.5 |
| MolProbity score | 2.19 | 2.13 | 1.95 | 1.42 |
| Bad bonds (%) | 0.03 | 0 | 0 | 0 |
| Bad angeles (%) | 0.06 | 0.03 | 0.02 | 0 |

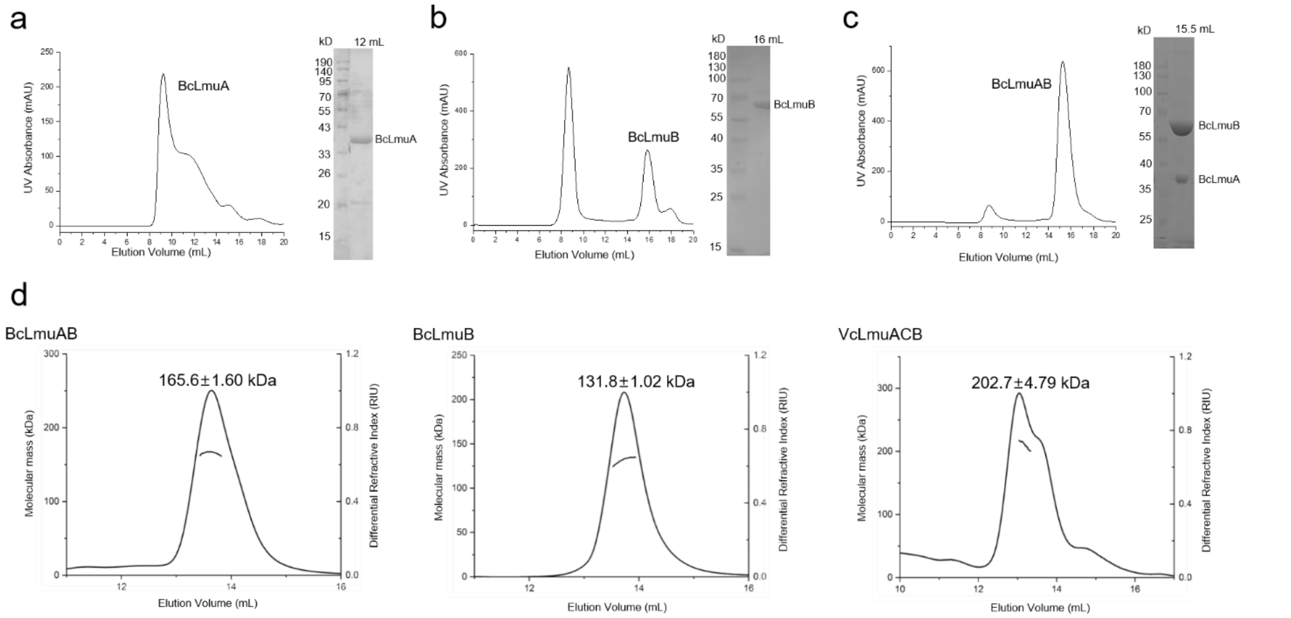


e

| Protein | Actual Molecular Weight (kDa) | Assume Ratio (A:B:C) | Theory Molecular Weight (kDa) |
|---|---|---|---|
| *Bc*LmuAB | 165.6±1.60 | 1:2 | 168.90 |
| *Bc*LmuB | 131.8±1.02 | 2 | 132.84 |
| *Vc*LmuACB | 202.7±4.79 | 1:2:1 | 213.09 |

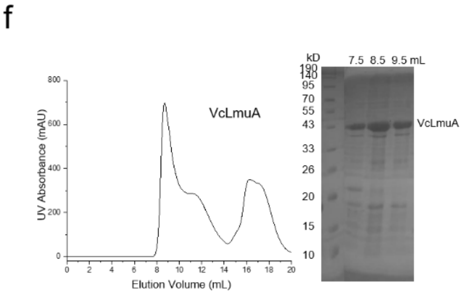


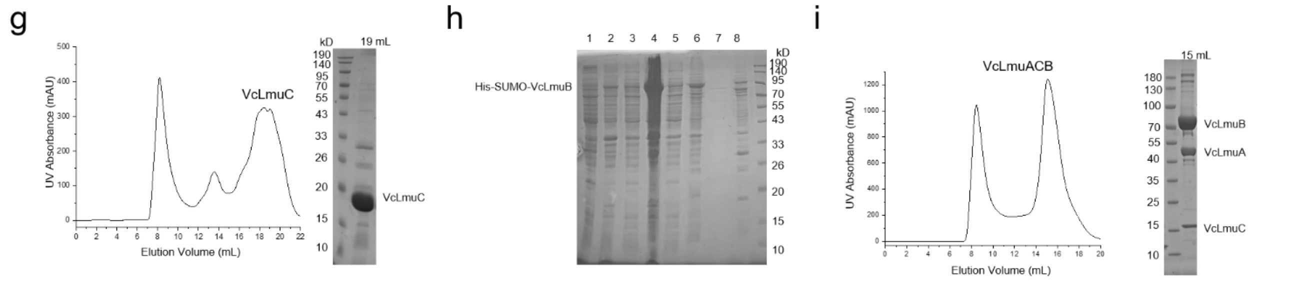


**Extended Data Fig.1|Type I and type II Lamassu systems both form heterooligomers**

**a-c**, Purification profile of BcLmuA (a), BcLmuB (c) and BcLmuAB complex (c) on Superdex 200 column.

**d**,SEC-MALS (Size Exclusion Chromatography with Multi-Angle Light Scattering) analysis of BcLmuB, BcLmuAB and VcLmuACB complex. Respective calculated molecular weight is marked on top of the peak.

**e**,Statistics of the SEC-MALS results and theoretical molecular weight based on predicted complex ratios.

**f-g**, Purification profile of VcLmuA (f) and VcLmuC (g) on Superdex-200 column.

**h**, Purification test of His-SUMO-TEV-VcLmuB and SDS-PAGE gel of the samples. 1-bacteria pre-induced. 2-bacteria post-induced. 3-supernatant. 4-pellet. 5-flowthrough after Ni-column. 6-beads after wash. 7-Elution. 8-beads after elution.

**i**, Purification profile of VcLmuACB complex on Superdex 200 column.

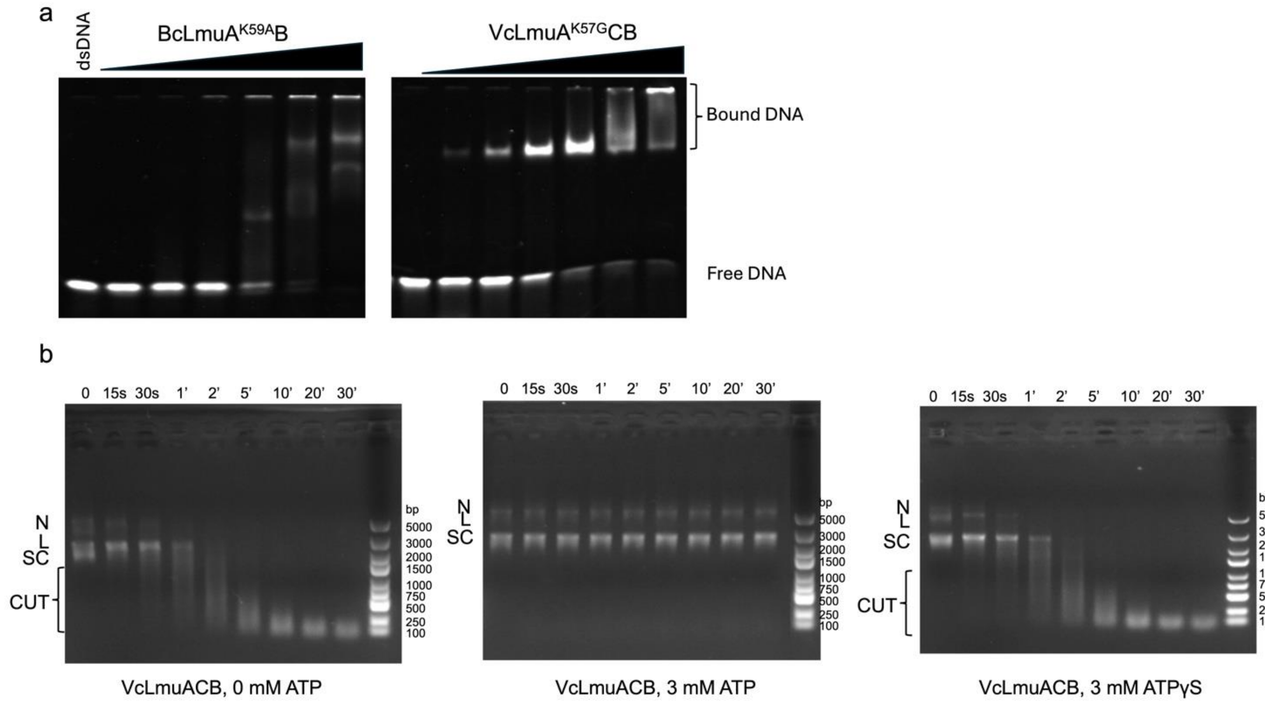


**Extended Data Fig.2|DNA binding and nuclease activity of the Lamassu system**

**a**, EMSA to test the binding of BcLmuA$^{K59A}$B and VcLmuA$^{K57G}$CB to dsDNA. BcLmuA$^{K59A}$B and VcLmuA$^{K57G}$CB is included at a concentration of 0.25, 0.5, 1, 2, 4, 8 μM, respectively.

**b**, pUC19 digestion by VcLmuACB in the presence or absence of ATP or ATPγS, respectively.

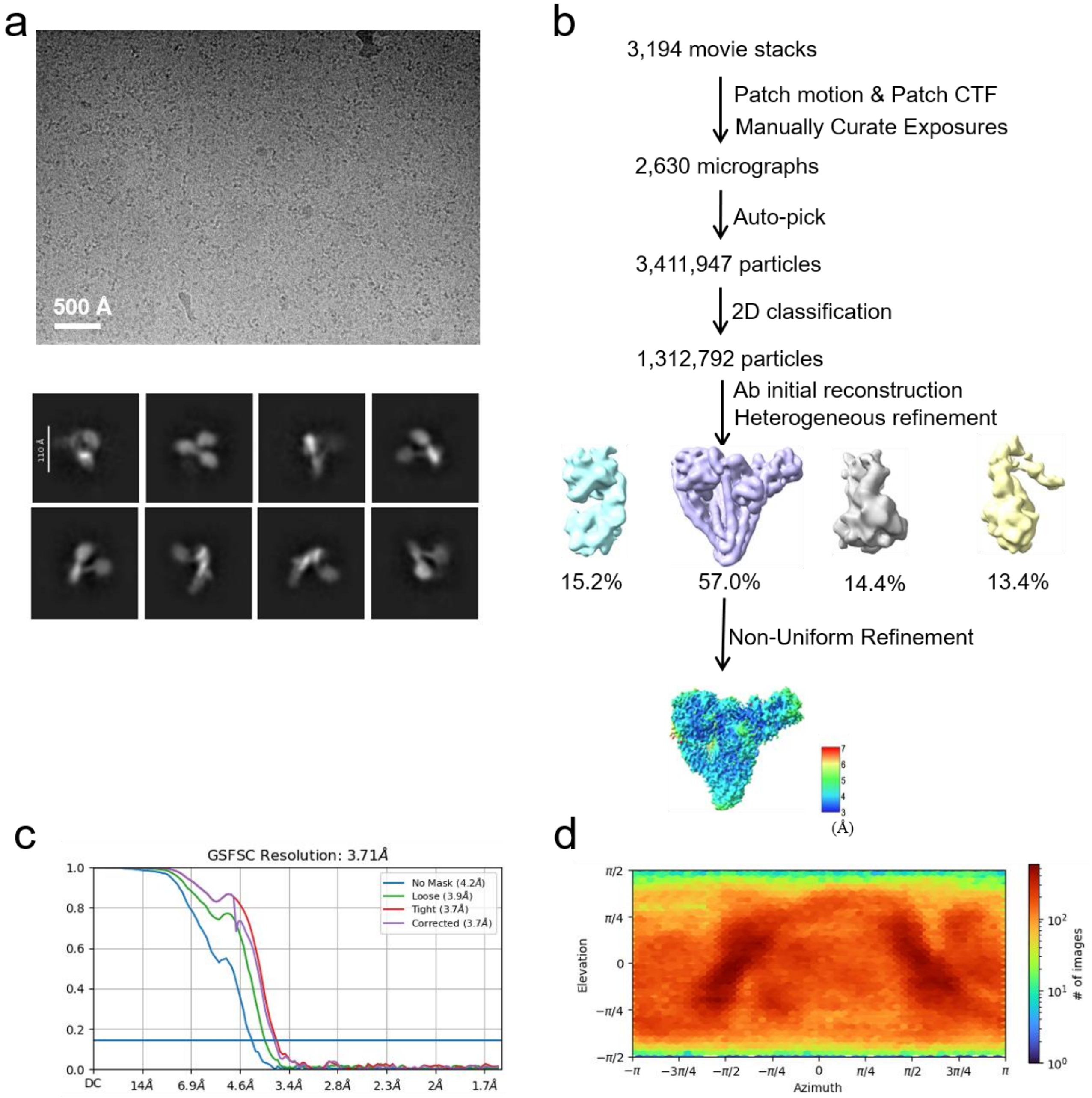


**Extended Data Fig.3|Single-particle cryo-EM analysis of the BcLmuAB complex.**

**a**,Representative cryo-EM micrograph (upper) and reference-free 2D class averages (lower) of BcLmuAB.

**b**,Workflow of the cryo-EM data processing. The final map resolution is color coded for different regions.

**c**,Gold standard FSC plots for the 3D reconstructions of the whole map, calculated in cryoSPARC.

**d**,Euler angle distribution of the particle images.

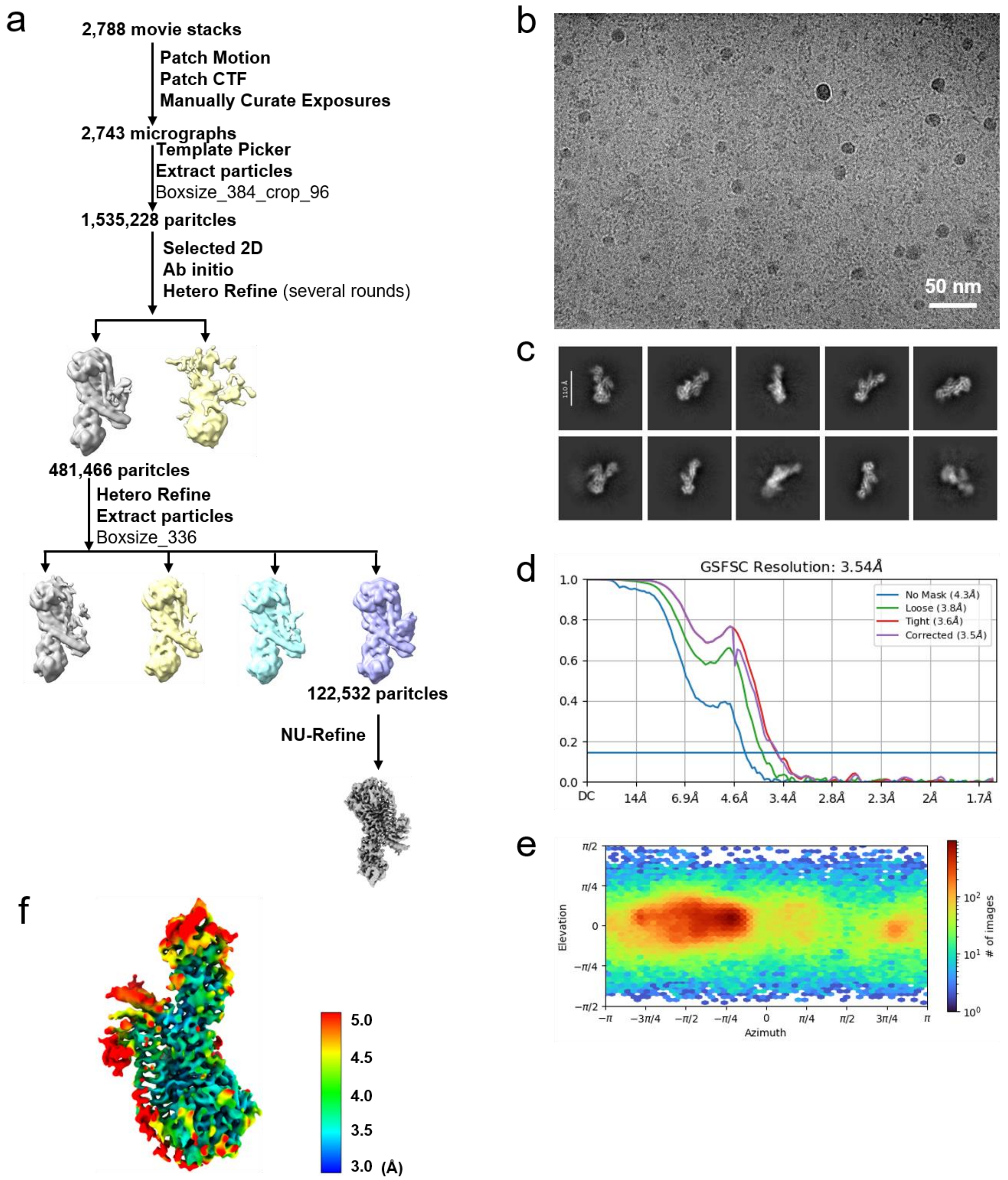


**Extended Data Fig.4|Single-particle cryo-EM data processing of VcLmuACB complex.**

**a**,Workflow of cryo-EM data processing of VcLmuACB.

**b**,Representative cryo-EM micrograph of VcLmuACB.

**c**,Reference-free 2D class averages of VcLmuACB.

**d**,Gold standard FSC plot for the final 3D reconstruction of the whole map, calculated in cryoSPARC.

**e**,Euler angle distribution of the particle images.

**f**,The final map resolution is colored for different regions.

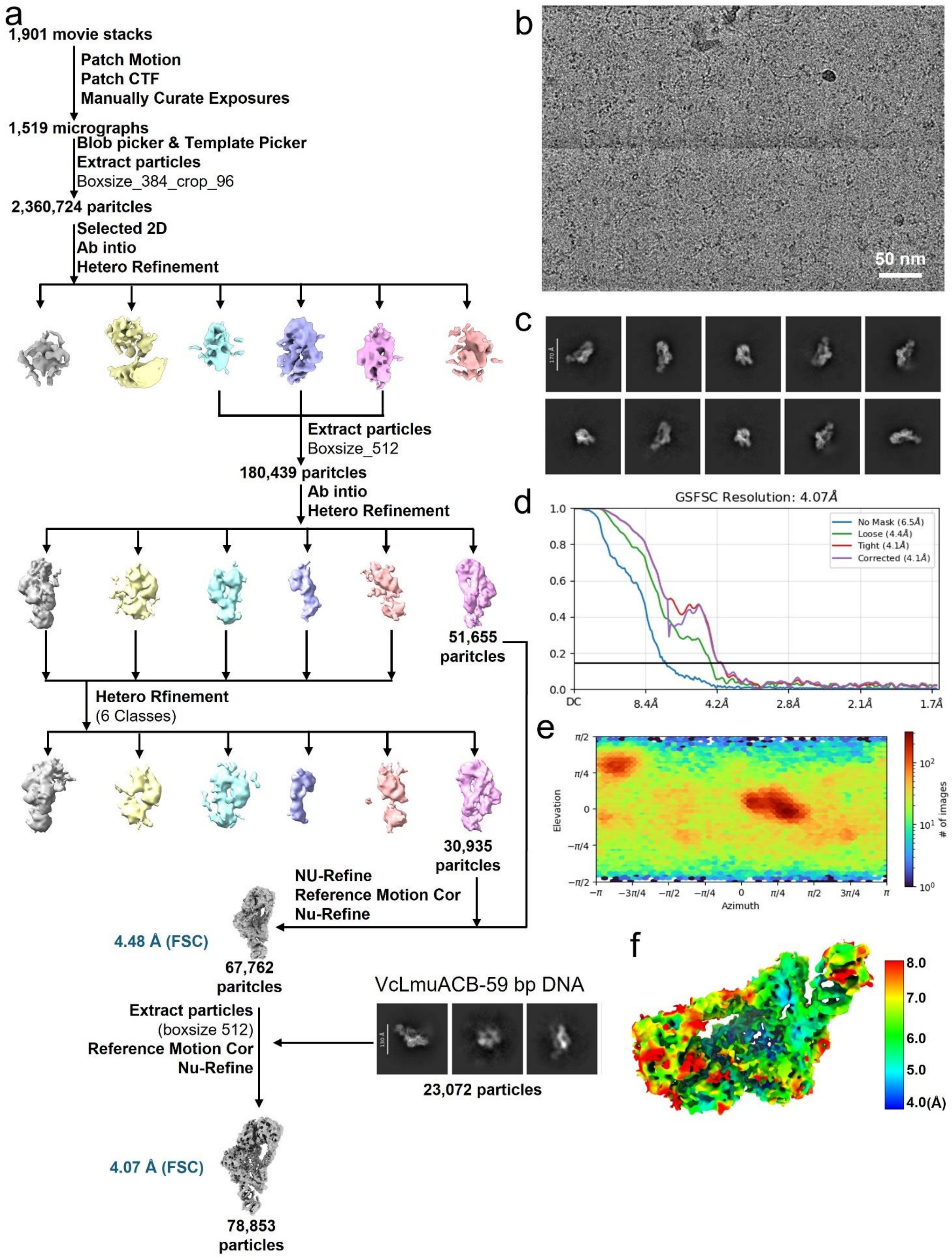


**Extended Data Fig.5|Single-particle cryo-EM data processing of VcLmuACB-pUC19.**
**a**,Workflow of cryo-EM data processing of VcLmuACB-pUC19.

**b**,Representative cryo-EM micrograph of VcLmuACB-pUC19.
**c**,Reference-free 2D class averages of VcLmuACB-pUC19.
**d**,Gold standard FSC plot for the final 3D reconstruction of the whole map, calculated in cryoSPARC.
**e**,Euler angle distribution of the particle images.
**f**,The final map resolution is colored for different regions.

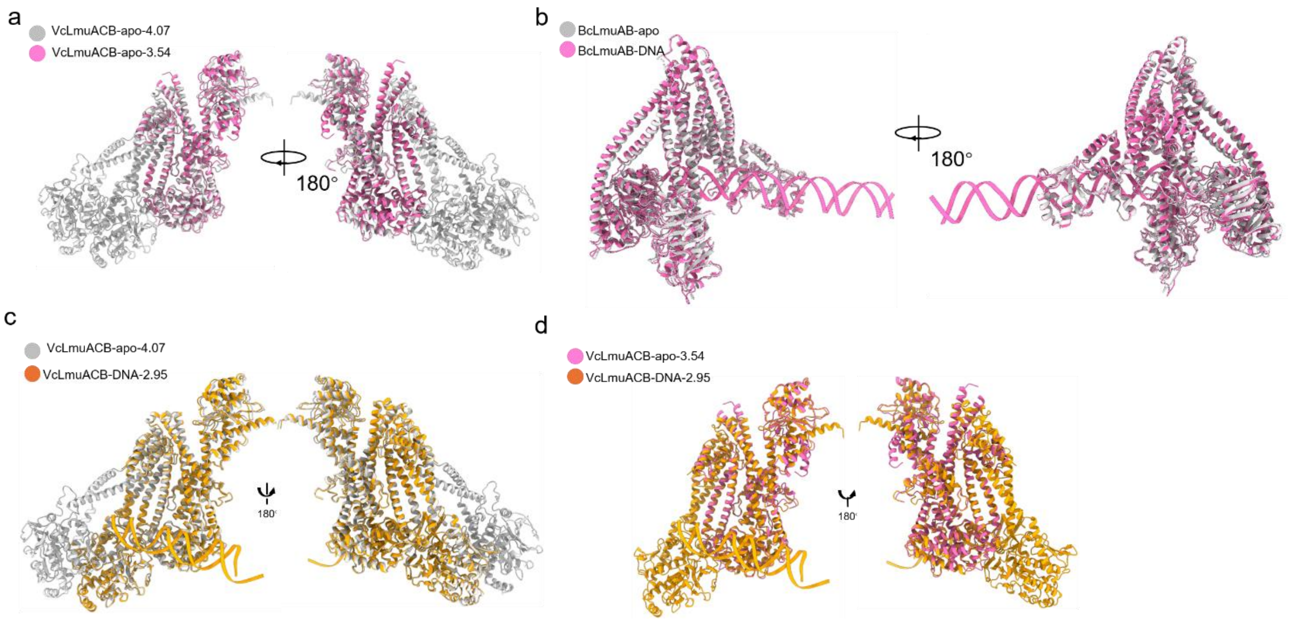


**Extended Data Fig.6|Structural alignment between different states of BcLmuAB and VcLmuACB complexes.**

**a**,Structural alignment between the two apo state structures of VcLmuACB with different protein lengths.

**b-d**,Structural alignment between the apo and DNA-bound states of BcLmuAB (b), between the apo (partial structure) and DNA-bound states of VcLmuACB (c), and between the apo (full length) and DNA-bound states of VcLmuACB (d).

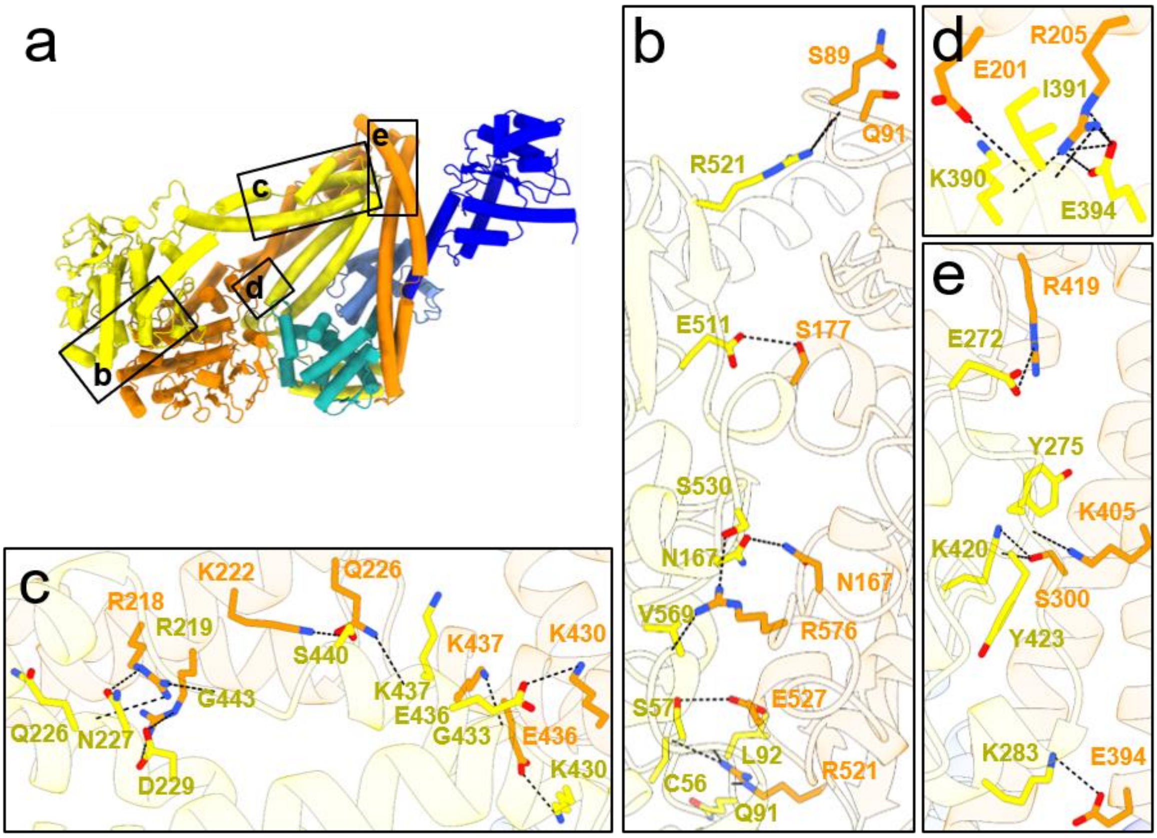


**Extended Data Fig.7|Inter-subunit interactions in the apo state of VcLmuACB (full length).**
**a-e**,Atomic model of the VcLmuACB complex is shown in a, in which detailed interactions between dsDNA and BcLmuAB are shown in b-e.

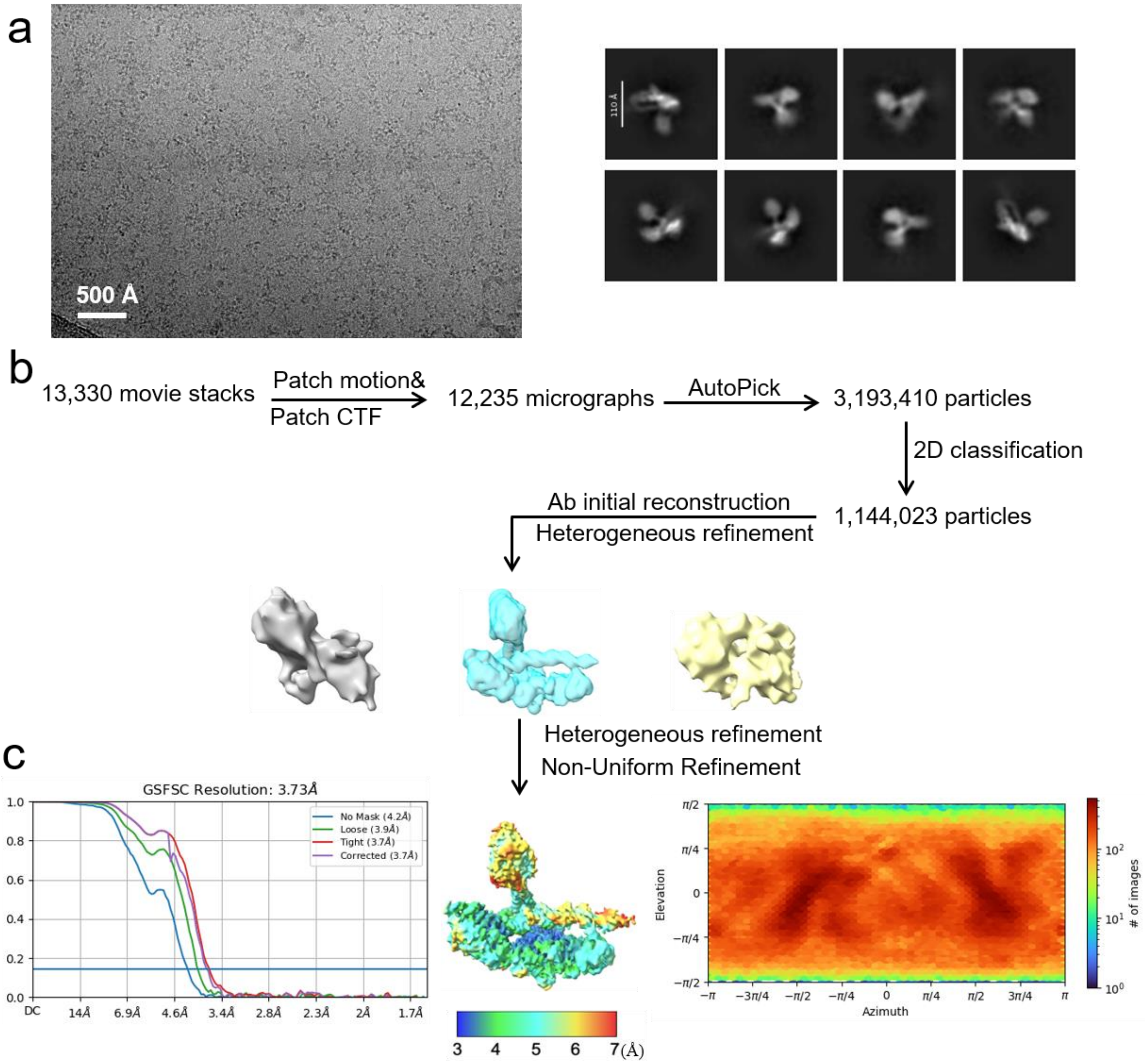


**Extended Data Fig.8|Single-particle cryo-EM analysis of the BcLmuAB-DNA complex.**

**a**,Representative cryo-EM micrograph (left) and reference-free 2D class averages (right) of BcLmuAB-DNA.

**b**,Workflow of the cryo-EM data processing. The final map resolution is color coded for different regions.

**c**,Gold standard FSC plots for the 3D reconstructions of the whole map, calculated in cryoSPARC (left) and Euler angle distribution of the particle image(right).

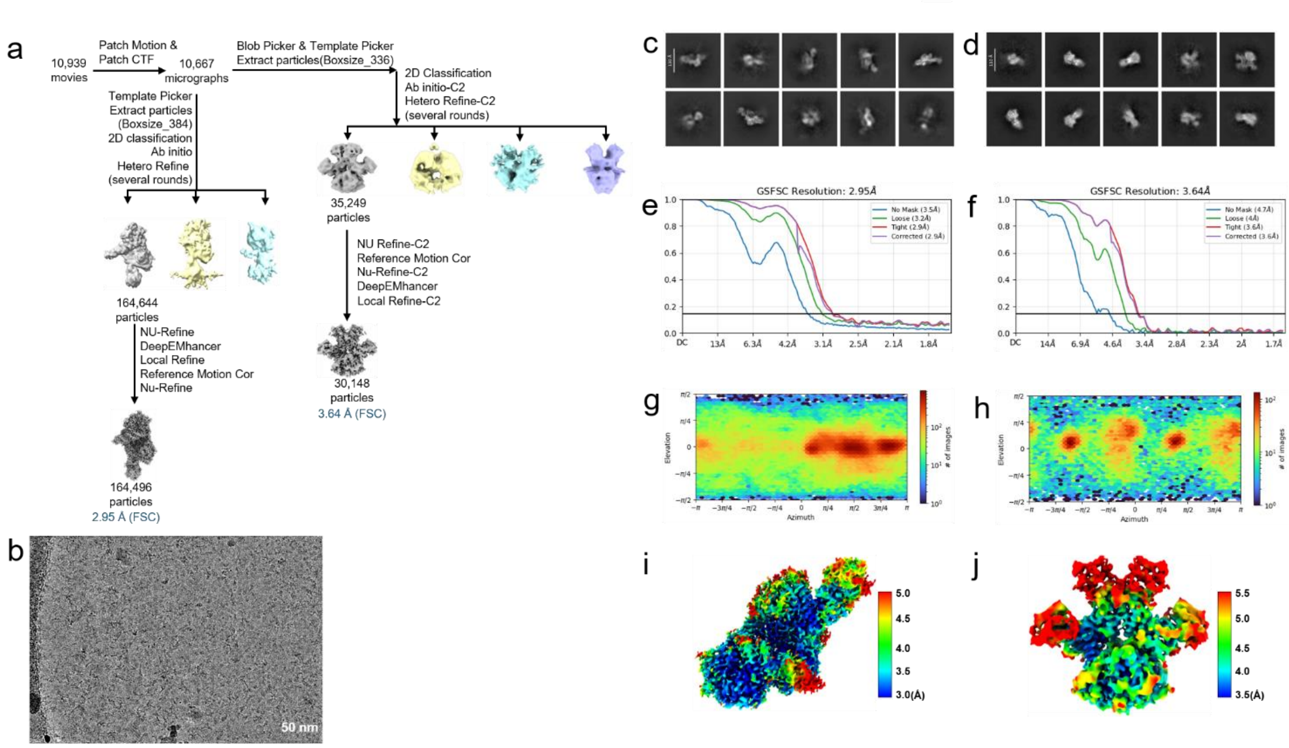


**Extended Data Fig.9|Single-particle cryo-EM data processing of VcLmuACB-59bp DNA complex and VcLmuA-tetramer complex.**

**a**,Workflow of cryo-EM data processing of VcLmuACB-59bp DNA.

**b**,Representative cryo-EM micrograph of VcLmuACB-59bp DNA.

**c**,Reference-free 2D class average of VcLmuACB-DNA.

**d**,Reference-free 2D class average of VcLmuA-tetramer.

**e**,Gold standard FSC plots for the 3D reconstructions of the whole map of VcLmuACB-DNA, calculated in cryoSPARC.

**f**,Gold standard FSC plots for the 3D reconstructions of the whole map of VcLmuA-tetramer, calculated in cryoSPARC.

**g**,Euler angle distribution of the particle image of VcLmuACB-DNA.

**h**,Euler angle distribution of the particle image of VcLmuA-tetramer.

**i,**The final map resolution is color coded for different regions of VcLmuACB-DNA.

**j**,The final map resolution is color coded for different regions of VcLmuA-tetramer.

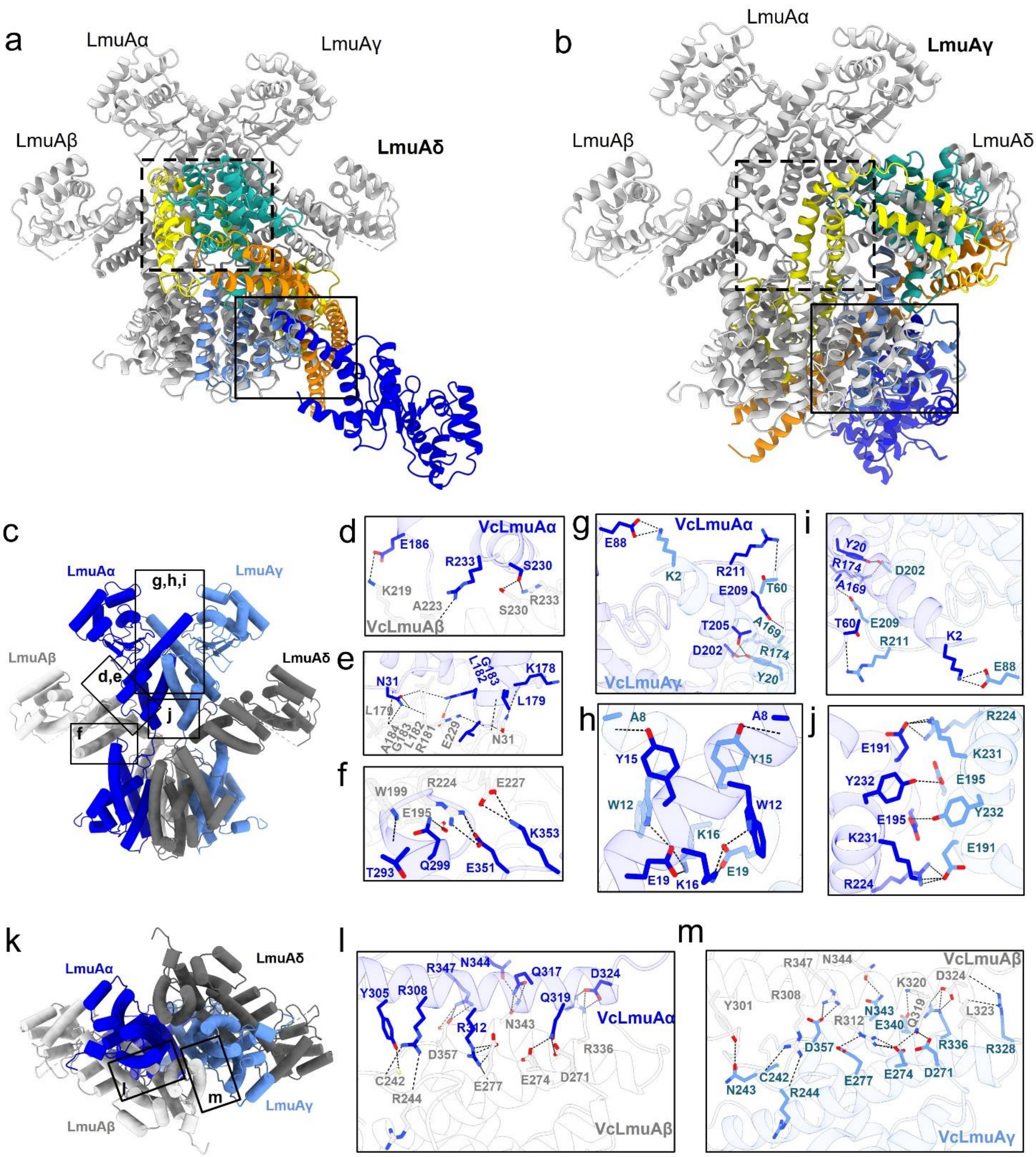


**Extended Data Fig.10|Detailed interactions among the subunits of the VcLmuA tetramer**

**a-b**,Structural alignment between VcLmuA tetramer and VcLmuACB complex (the one with partial LmuB structure) at VcLmuAδ-CTD (a) and VcLmuAγ-CTD (b), respectively. The VcLmuACB complex is colored as in Fig.2b, and the VcLmuA tetramer is colored gray. The superimposed VcLmuA protomer is marked in bolded form.

**c-j**,Atomic model of the VcLmuA tetramer (sideview) is shown in a, in which detailed interactions between NTDs and between NTDs and CTDs are shown in B-H.

**k-m**,Atomic model of the VcLmuA tetramer (bottom view) is shown in i, in which detailed interactions between CTDs are shown in j and k.

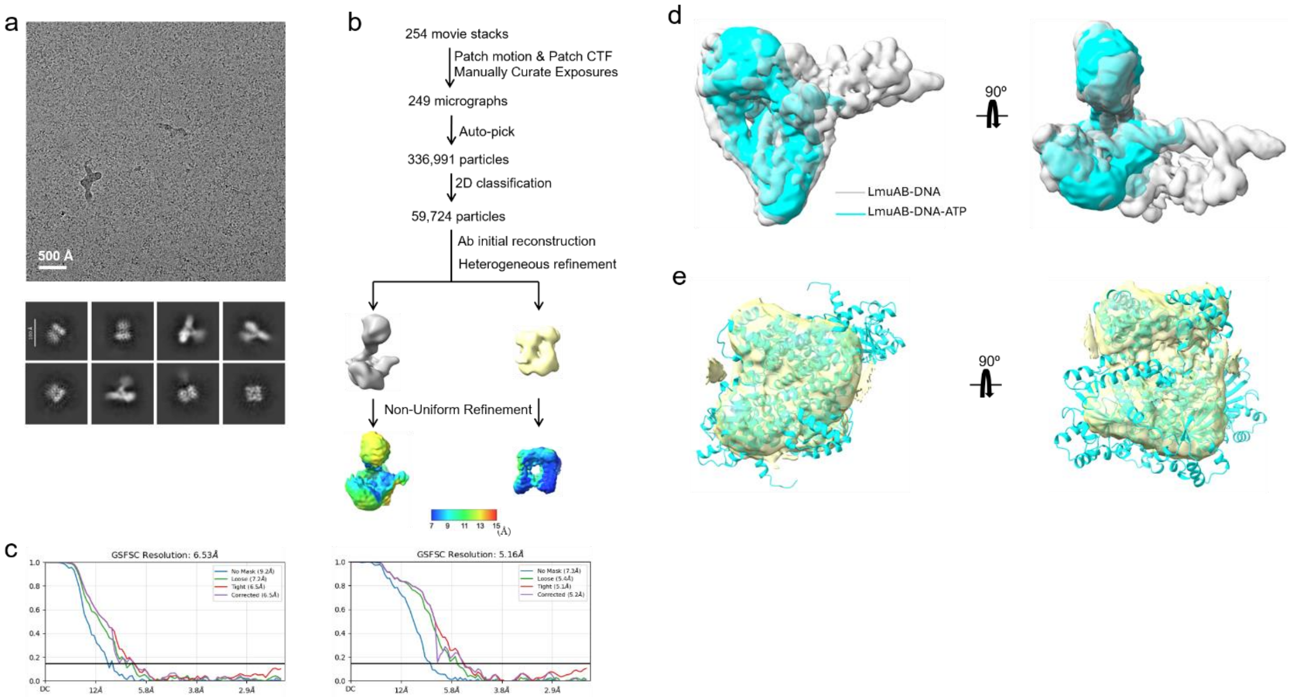


**Extended Data Fig.11|Single-particle cryo-EM analysis of the BcLmuA(K59A)B-DNA-ATP complex.**

**a**,Representative motion-corrected cryo-EM micrograph (upper) and reference-free 2D class averages (lower) of LmuAB(K59A)-DNA-ATP.

**b**,Workflow of the cryo-EM data processing. The final map resolution is color coded for different regions.

**c**,Gold standard FSC plots for the 3D reconstructions of the whole map, calculated in cryoSPARC.

**d**,Overlay of the cryo-EM maps of the LmuAB–DNA complex (gray) and the LmuB–DNA complex (light sea green) for structural comparison.

**e**,Predicted model of the LmuA tetramer fitted into the cryo-EM density map.

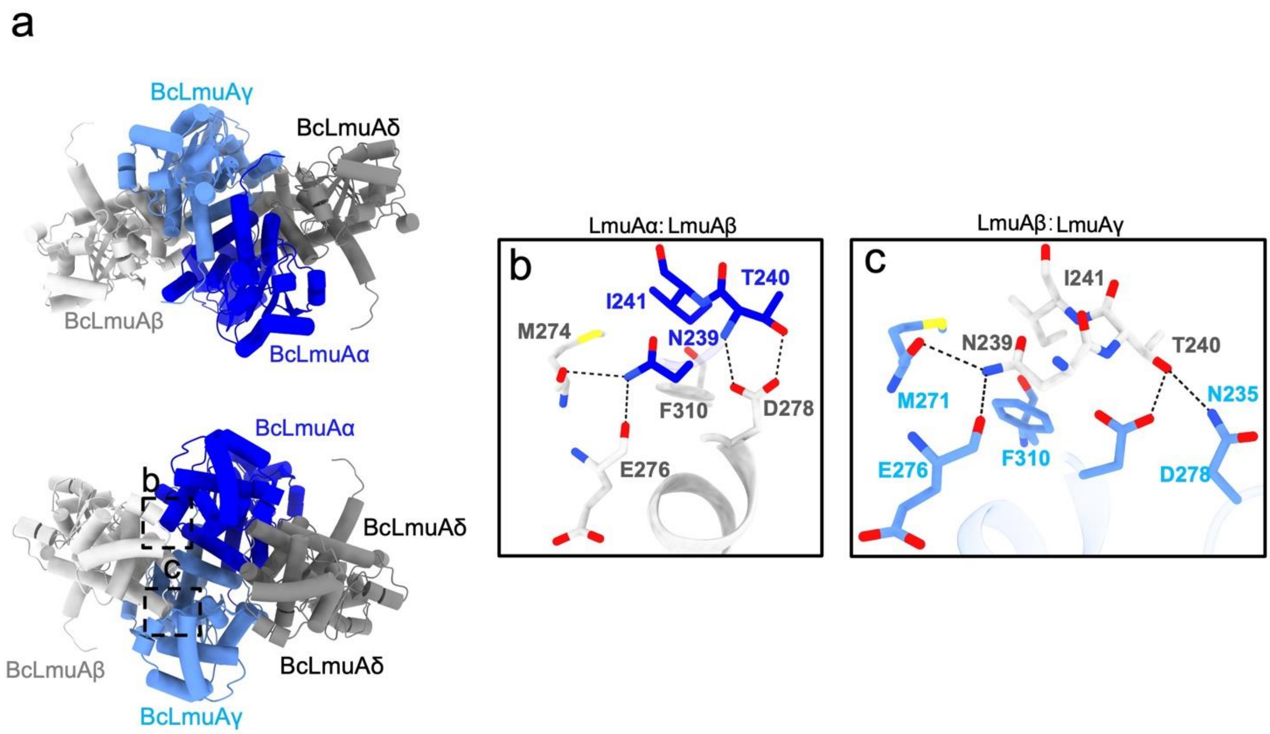


**Extended Data Fig.12| Predicted structural model of BcLmuA tetramer.**

**a**,Predicted structural model of BcLmuA tetramer.

**b-c**,Detailed interactions among the subunits of the predicted BcLmuA tetramer.
